# The *C. elegans* Connectome Toolbox: an open, extensible framework for multimodal connectomic data integration and analysis

**DOI:** 10.64898/2026.11.04.764366

**Authors:** Yasinthan Vickneswaran, Adam Ponzi, Ankur Sinha, Netta Cohen, Stephen D. Larson, Padraig Gleeson

## Abstract

The nematode *C. elegans* is one of the best studied model organisms in biology. Since the initial description of the *C. elegans* neuronal connectome by White and colleagues, numerous studies have mapped and analyzed the connectome, and released structured datasets of the connectivity, often merging and adding to previous versions. These datasets now serve as foundational resources that underpin studies into the neuronal basis of *C. elegans* behavior. In spite of their high utility, heterogeneity in the different datasets and their annotation conventions obscures direct comparison where discrepancies between connectomes frequently stem from differences in coverage or nomenclature rather than genuine biological disagreement. Additionally, datasets describing new connectivity modalities - contact-based (the contactome), extrasynaptic (monoaminergic and neuropeptidergic), and functional connectivity - have also recently become available. Thus, there is a need for a unified framework providing standardized access to these diverse datasets while preserving their individual provenance. Here, we present the *C. elegans* Connectome Toolbox, a comprehensive framework for consolidating, curating and validating historical and newly published multimodal *C. elegans* connectivity data. Connectome Toolbox includes an accessible online resource (https://openworm.org/ConnectomeToolbox) for interactive data exploration and is accompanied by a Python library for programmatic access. We demonstrate the utility of the toolbox through a series of analyses that illustrate how connectome data can be explored, visualized, and compared across modalities, including integration of the anatomical connectome with a comprehensive neurotransmitter atlas, and comparison with computational models of *C. elegans* neural circuits.

## Introduction

Understanding the neuronal basis of behavior remains a central challenge in neuroscience. Behavior arises as an emergent property of the connectivity among biological neurons within the context of the animal’s embodiment and situatedness (***Brooks, 2018***). Elucidating the wiring of the nervous system is therefore essential for interpreting the complexity of behavior (***Swanson and Lichtman, 2016***; ***Emmons, 2024***). Volume Electron Microscopy (EM) and automated reconstruction have transformed the scale at which this is possible, from local circuits (***Lichtman and Sanes, 2008***) to whole-brain wiring diagrams (***MICrONS Consortium, 2025***; ***Dorkenwald et al., 2024***), yet anatomical connectivity alone does not capture all modes of neuronal communication (***Marder and Thirumalai, 2002***; ***Bargmann, 2012***). Neurons also interact via extrasynaptic, “wireless” signaling pathways (***Bentley et al., 2016***; ***Ripoll-Sánchez et al., 2023***), which can have a critical but often ignored influence on the effective functional connectivity within neural networks (***Randi et al., 2023***). A mechanistic understanding of behavior therefore requires not just mapping these individual connectivity modalities, but integrating them.

The nematode *C. elegans* is currently the only organism where this integration is tractable across scales, from the nanoscopic to whole brain to behavior, and its advantages are well established. Its ease of cultivation, short life cycle, and compact genome have made it exceptionally amenable to experimental study (***Corsi et al., 2015***). Responding to Sydney Brenner’s founding proposal (***Brenner, 1974***), the community systematically resolved the complete cell lineage from zygote to adult (***Sulston and Horvitz, 1977***; ***Sulston et al., 1983***), produced the first fully sequenced genome of a multicellular organism (***The*** C. elegans ***Sequencing Consortium, 1998***), and generated the first comprehensive reconstruction of an entire nervous system (***White et al., 1986***). Its synaptic and gap-junction connectome has been reconstructed at cellular resolution across sexes and developmental stages (***White et al., 1986***; ***Cook et al., 2019***; ***Witvliet et al., 2021***); its monoaminergic and neuropeptidergic signaling networks have been mapped across the entire nervous system from gene-expression and receptor-binding data (***Bentley et al., 2016***; ***Ripoll-Sánchez et al., 2023***); and its neurotransmitter usage has been cataloged neuron by neuron (***Taylor et al., 2021***; ***Wang et al., 2024***). Every neuron has a stereotyped genetic identity and a known position, so molecular and anatomical layers can be overlaid directly onto the same structural scaffold. Together they have enabled whole-brain calcium imaging of behavior in freely moving animals (***Atanas et al., 2023***) and systematic optogenetic perturbation of identified cells (***Randi et al., 2023***). No other organism currently offers this combination of completeness across scales and modalities.

Before describing the framework we have built to facilitate access to this data, it is useful to define the principal categories of connectivity information now available for *C. elegans*:

1. The anatomical connectome: which neurons are connected via chemical and electrical (gap junction) synaptic connections (***Albertson and Thomson, 1976***; ***White et al., 1986***; ***Durbin, 1987***; ***Hall and Russell, 1991***; ***Varshney et al., 2011***; ***Jarrell et al., 2012***; ***Sammut et al., 2015***; ***Cook et al., 2019***, ***2020***; ***Molina-García et al., 2020***; ***Witvliet et al., 2021***)?
2. The contactome: which neurons are in direct physical contact, forming membrane appositions (***Brittin et al., 2021***; ***Moyle et al., 2021***; ***Witvliet et al., 2021***; ***Yim et al., 2024***)?
3. Neurotransmitter atlases: which neurotransmitters are released by each neuron (*Serrano-Saiz et al., 2013*; *Pereira et al., 2015*; *Gendrel et al., 2016*; *Taylor et al., 2021*; *Beets et al., 2023*; *Dag et al., 2023*; *Wang et al., 2024*), and are their outputs predominantly excitatory or inhibitory (*Fenyves et al., 2020*)?
4. The extrasynaptic connectome: based on gene-expression data, which neurons release and which possess receptors for specific monoamines and neuropeptides (***Bentley et al., 2016***; ***Ripoll-Sánchez et al., 2023***)?
5. The functional connectome: what is the sign, strength and directionality of each neuron’s causal influence on others, obtained by optogenetically stimulating one neuron and recording the response of others (***Randi et al., 2023***)? It should be noted that this differs from “functional connectivity” as used in neuroimaging, which refers to statistical correlation between regional activity.

The history of *C. elegans* connectomics reflects decades of progressive refinement across these modalities (Fig. 1). Brenner’s group pioneered the first anatomical reconstructions, beginning with the pharyngeal nervous system (***Albertson and Thomson, 1976***), and the ventral nerve cord (***White et al., 1976***), before White and colleagues produced the first complete reconstruction of an entire nervous system in any organism (***White et al., 1986***). That seminal study identified 302 neurons grouped into 118 classes (later confirmed genetically (***Reilly et al., 2020***)) and cataloged approximately 5,000 chemical synapses, 2,000 neuromuscular junctions (NMJs), and 600 gap junctions. Durbin subsequently organized this synaptic connectivity data into a digital database (***Durbin, 1987***), transforming the connectome from a static anatomical description into a computationally tractable resource and laying the foundation for connectomics as a discipline.

**Figure 1.**
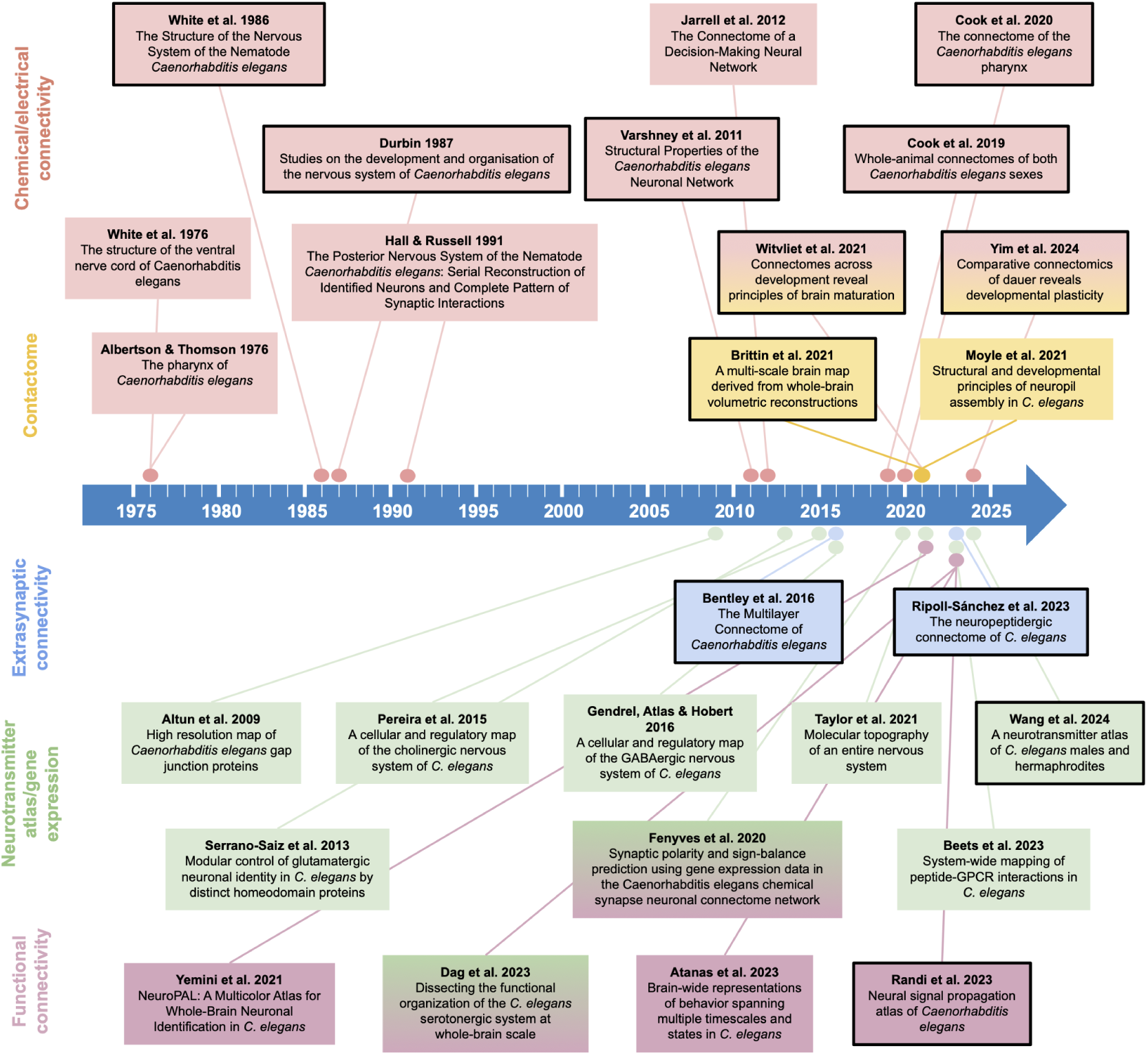
An overview of published studies on *C. elegans* neuronal connectivity and synaptic transmission. Data has been published on the anatomical connectivity (both numbers of identified chemical and electrical synapses and the contactome – locations of physical contact between cells), the extrasynaptic connectome, gene expression studies identifying which neurotransmitters are released and which receptors are present in identified neurons, and on the functional connectivity of neurons. Papers highlighted with a black border have released structured datasets on connectivity which have been incorporated into the *C. elegans* Connectome Toolbox.

Building on these foundational studies, Chen and colleagues used a newly compiled version of the wiring diagram to test Cajal’s wiring economy principle at the resolution of individual neurons, solving for the cost-minimizing layout of the 279 non-pharyngeal neurons (***Chen et al., 2006***). Most neurons lay close to their predicted positions, while those that deviated most strongly proved to share stereotyped roles, so that the departures from optimality were themselves informative about function. This was an important early demonstration of what becomes possible once connectivity is available as structured, computable data. They subsequently produced a comprehensive, self-consistent connectome of the somatic nervous system, incorporating the connectivity of the posterior nervous system (***Hall and Russell, 1991***) through reanalysis of the original EM material, and performed quantitative analyses of its statistical and topological properties (***Varshney et al., 2011***).

Until this point, most reconstructions focused on the adult hermaphrodite. The male posterior nervous system was addressed by Jarrell and colleagues (***Jarrell et al., 2012***) and subsequently refined with additional information on neuronal identity and connectivity (***Sammut et al., 2015***; ***Molina-García et al., 2020***). Comprehensive connectomes of both sexes were eventually presented by Cook and colleagues (***Cook et al., 2019***), providing quantitative connectivity matrices spanning sensory neurons to muscles and other target cells such as non-muscle end organs. The pharyngeal wiring diagram was later revisited at individual neuron resolution, revealing a shallow circuit architecture in which most neurons possess sensory endings and all connect directly to end organs (***Cook et al., 2020***).

While the connectome maps synaptic and electrical connections between neurons, physical contact between neurites is far more extensive than what is captured by these functional specializations alone. Brittin and colleagues introduced the contactome - a comprehensive, nanoscale-resolution mapping of all membrane contacts between neurons from whole-brain volumetric reconstructions (***Brittin et al., 2021***). The contactome provides the spatial scaffold needed to characterize the modular organization of the nerve ring, identify candidate sites of non-synaptic interaction, and quantify inter-individual variability in neural architecture. By quantifying spatial proximity independent of synaptic specialization, the contactome revealed the extent to which neurite adjacency both constrains and exceeds synaptic connectivity. Moyle and colleagues further analyzed these contact relationships using alternative clustering approaches to generate adjacency representations of neuronal proximity within the nerve ring (***Moyle et al., 2021***).

More recently, connectomes have been reconstructed at eight developmental time points from birth to adulthood, revealing that connection number, path length, and clustering all follow systematic developmental trajectories (***Witvliet et al., 2021***). The dauer-stage connectome has further captured structural reorganization associated with this stress-induced state (***Yim et al., 2024***). Both of these studies have also analyzed the worm’s contactome.

In parallel with these studies on anatomical connectivity, the extrasynaptic landscape has been mapped through the monoaminergic connectome (***Bentley et al., 2016***), and a comprehensive neuropeptidergic connectome generated using large-scale GPCR deorphanization and single-neuron scRNA-seq (***Ripoll-Sánchez et al., 2023***). Neurotransmitter usage across the nervous system has been cataloged in detail (***Taylor et al., 2021***; ***Wang et al., 2024***) and functional connectivity has been mapped using optogenetic stimulation and calcium imaging at a whole-brain scale (***Randi et al., 2023***).

Each of these categories of connectivity information has been addressed by multiple independent studies, each study introducing its own scope, annotation conventions, and file formats. The consequences of this fragmentation are concrete: direct comparison across datasets is non-trivial, and apparent biological differences between connectomes sometimes reflect differences in coverage or nomenclature rather than genuine disagreement. Even the anatomical boundaries of what constitutes the *C. elegans* connectome differ between studies. The connectomes published in ***Cook et al. (2019)***, for instance, include neurons alongside neuronal support cells, marginal cells, and muscles, whereas earlier reconstructions did not. Connectivity data for a single neuron are often distributed across multiple publications in heterogeneous formats, making integrated analysis laborious and error-prone. While datasets should ideally be released in structured formats as supplementary information associated with publications, in practice they are often released, refined or corrected post publication. Despite the existence of valuable community resources including WormAtlas (***Altun et al., 2025***), WormBase (***Harris et al., 2020***), WormWiring (***Cook et al., 2019***), and NemaNode (***Witvliet et al., 2021***), these platforms remain siloed; each exposes a different subset of data through a different interface, and a common, standardized method for programmatic access to documented, curated datasets, cross-dataset comparison, and direct integration with simulation frameworks is missing.

To address this fragmentation and make *C. elegans* connectomics datasets more Findable, Accessible, Interoperable, and Reusable (FAIR, ***Wilkinson et al. (2016)***), we developed a unified Python-based framework: the *C. elegans* Connectome Toolbox. The Toolbox rests on three design principles. First, multimodality: anatomical, contact-based, extrasynaptic, neurotransmitter, functional and model-derived connectivity are represented in a common internal, machine-readable format, so the same neuron or connection can be examined across modalities rather than studied in isolation. Second, standardization: all cells and groupings are mapped onto a common naming and annotation convention, so differences between datasets reflect the biology or methodology rather than nomenclature. Third, computational interoperability: the same machine-readable representations that drive the web visualizations can be read and written by downstream tools, including statistical analysis pipelines (for example, for examining bilateral symmetry) or use in neural and neuromechanical simulations.

The framework is fully open source and designed for extensibility: new datasets can be easily incorporated by adding a new dataset reader that translates it to the internal format, after which all existing visualization and analysis methods apply automatically. The Toolbox is accessible at multiple levels of technical expertise, from an interactive web interface requiring no programming experience (https://openworm.org/ConnectomeToolbox) to a Python package and API (Application Programming Interface) for programmatic analysis (*cect*, https://pypi.org/project/cect/). This work is a core part of the OpenWorm project (***Sarma et al., 2018***; ***Szigeti et al., 2014***), which has the long-term goal of digitally reconstructing *C. elegans*, and is intended as a living resource that can evolve as new datasets and modalities emerge. By unifying anatomical, extrasynaptic, and functional connectivity within a single computationally tractable framework, the *C. elegans* Connectome Toolbox offers both an immediate practical resource for the worm community and a methodological model for how multimodal connectomic infrastructure should be built as the field scales toward larger and more complex nervous systems.

## Results

### Connectome Toolbox enables consolidation, validation, visualization and analysis of ***C. elegans*** connectome datasets

We developed an open-source Python framework, the *C. elegans* Connectome Toolbox that provides uniform, machine-readable access to published *C. elegans* connectomic datasets, enabling direct comparison across studies and modalities. The sections that follow describe the package’s core data structures and visualization tools, then present a series of worked analyses spanning anatomical, neurotransmitter, contact-based, extrasynaptic, and functional connectivity, as well as comparison of empirical connectivity data with networks used in computational models of locomotion.

Fig. 2 gives an overview of the Python pipeline. Datasets of structured connectivity information are usually made available in the supplementary information of associated publications in one of many formats (e.g. text files, Excel/Open Office spreadsheets, comma separated values (CSV files), JSON). We developed custom Python readers (DataReader classes) for each source data file, which translate the connectivity data into our standard ConnectomeDataset class. This step involves the standardization of the naming of the neurons/muscles/support cells. Throughout, cells are identified using the naming conventions of ***Cook et al. (2019)*** as a common reference, with neurons classified as sensory neurons, interneurons, or motorneurons following conventions in that publication (some minor deviations from this are noted in the curation/validation notes in ***Supplementary file 1***).

**Figure 2.**
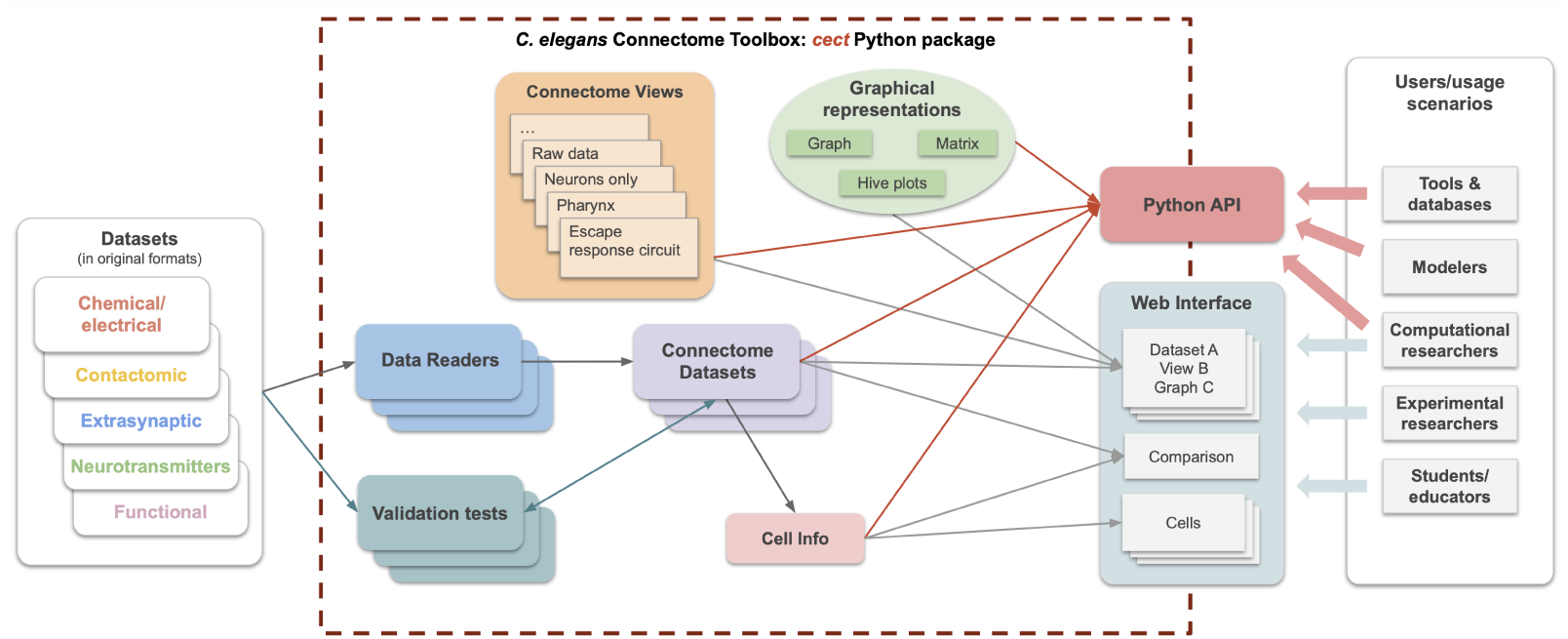
Python based pipeline for the *C. elegans* Connectome Toolbox Python package - *cect* (https://pypi.org/project/cect/). Each of the datasets of different modalities to be imported (left) have custom Python DataReader classes which convert connectivity data from their original formats (e.g. text files, Excel files, CSV files) into our standardized ConnectomeDataset class. Additionally, validation tests are written for each dataset, which enable automated checks that the imported data is consistent with the original published data. From the information in the ConnectomeDataset class, specific views of the data therein (e.g. all or a subset of cells; specific circuit views, specified in ConnectomeView classes) can be represented in multiple graphical visualization formats (e.g. adjacency matrices, connected graphs, hive plots) for exploration and analysis. A web interface for the package contains pregenerated graphical views for all permutations of these options, facilitating comparison between datasets. Dedicated pages for each cell summarize all known connectivity data for that cell, across all datasets and modalities. All of these functionalities are also available to users via the Python API (Application Programming Interface), enabling programmatic access to the data and visualizations for custom analyses. On the right, target users and example usage scenarios are listed, including use of the package for educational purposes, computational investigations of the neuronal basis of *C. elegans* behavior, and using the package as the basis for other tools and online resources. **Figure 2–Figure supplement 1**. Screenshot of a page on the Connectome Toolbox website for a single dataset. **Figure 2–Figure supplement 2**. Screenshot of a page on the website for a single cell. **Figure 2–Figure supplement 3**. Screenshot of a Jupyter notebook illustrating the use of the Python API.

Each ConnectomeDataset contains a list of the cells present in the specific dataset, as well as connections of different classes between them (e.g. independent adjacency matrices for chemical and electrical connections). These can be used directly by Python scripts, for example to access information on numbers of connections to/from individual cells of a particular type. For each imported dataset, we have also created a set of validation tests to verify that the data in the ConnectomeDataset is consistent with the original published data (e.g. total connection counts, specific weights of connections), along with documenting the sources of files and any errors/inconsistencies found (see Methods, ***Supplementary file 1***). Boxes highlighted with black borders in Fig. 1 indicate the publications on this timeline whose datasets have been incorporated, Fig. 3 shows the specific cells of different classes which are present in each, and Table 1 provides a complete list of datasets along with descriptions.

**Figure 3.**
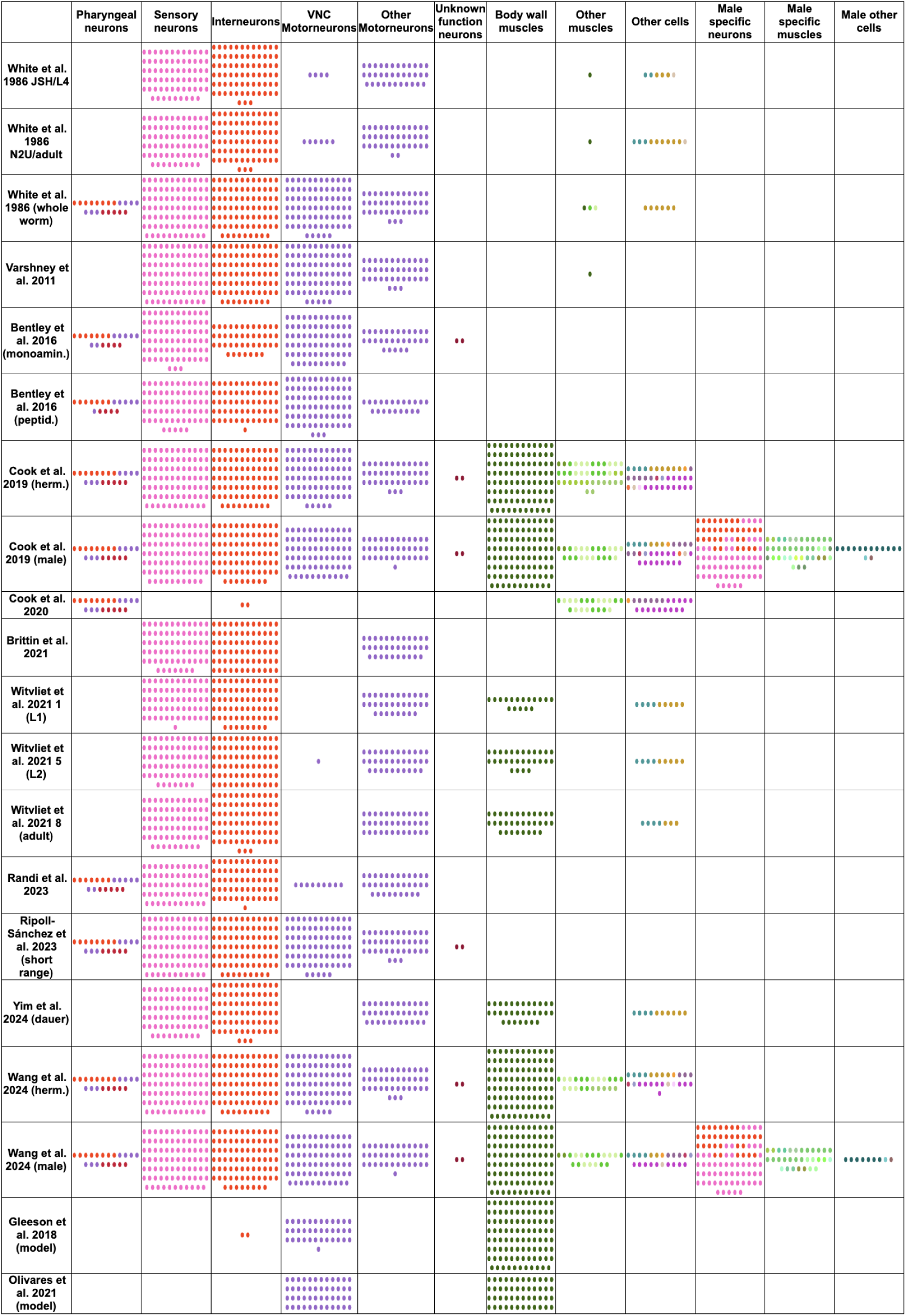
Numbers and types of cells present in datasets incorporated into the *C. elegans* Connectome Toolbox. See Table 1 for more details on each of these. Only 3 of 8 datasets from ***Witvliet et al. (2021)***, and one from each of ***Ripoll-Sánchez et al. (2023)*** and ***Yim et al. (2024)*** are included for brevity. Final two rows are cells present in computational models. Single points represent a single cell of that class. Colors based on WormAtlas colors (see Appendix 1 Fig. 1). An interactive version of this figure is at https://openworm.org/ConnectomeToolbox/Comparison. **Figure 3–Figure supplement 1.** Examples of graphical plots which can be generated for each connectome. **Figure 3–Figure supplement 2.** Comparison of numbers of connections to and from neurons across datasets. **Figure 3–Figure supplement 3.** Analysis of bilateral symmetry across datasets.

**Table 1.**
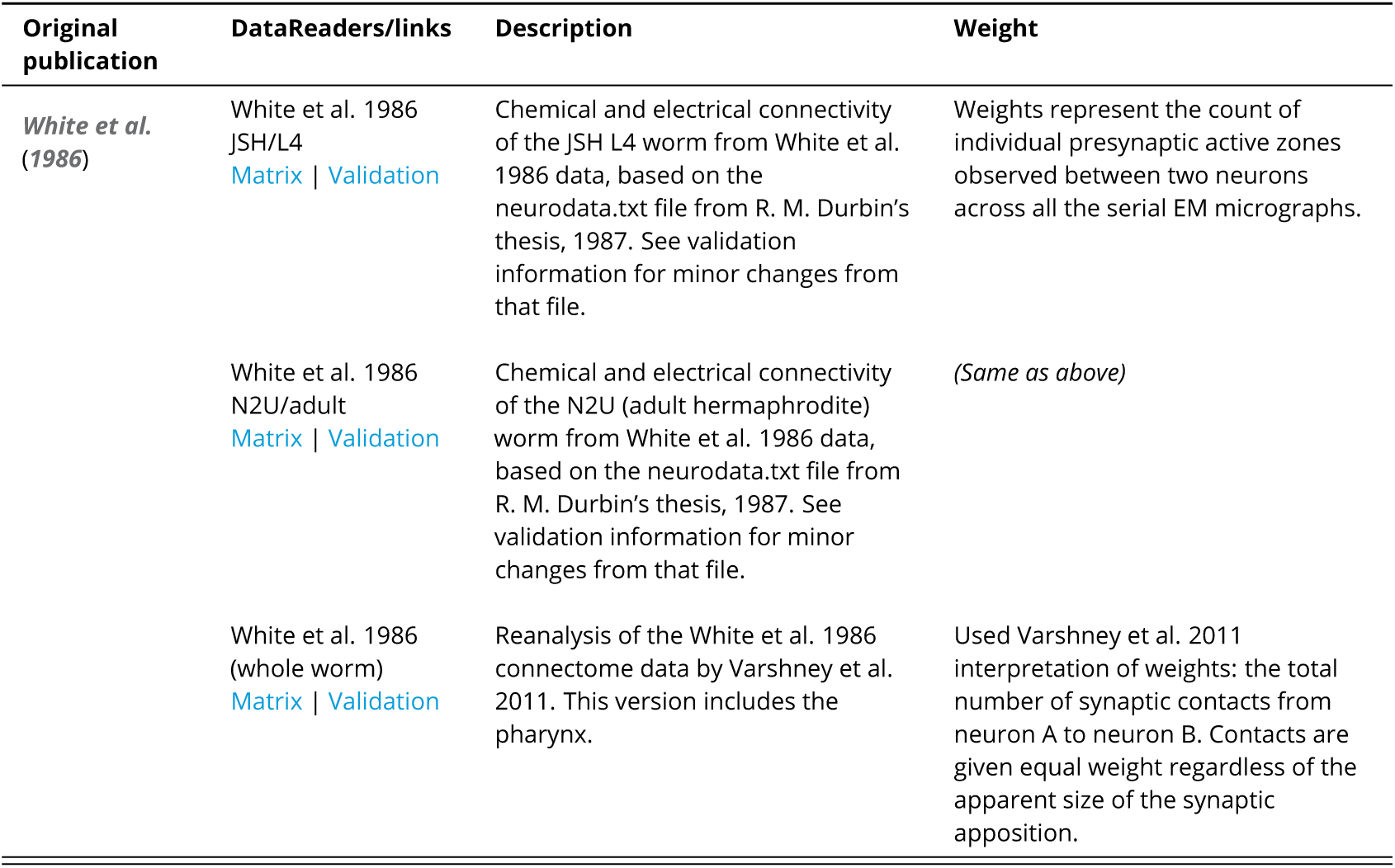
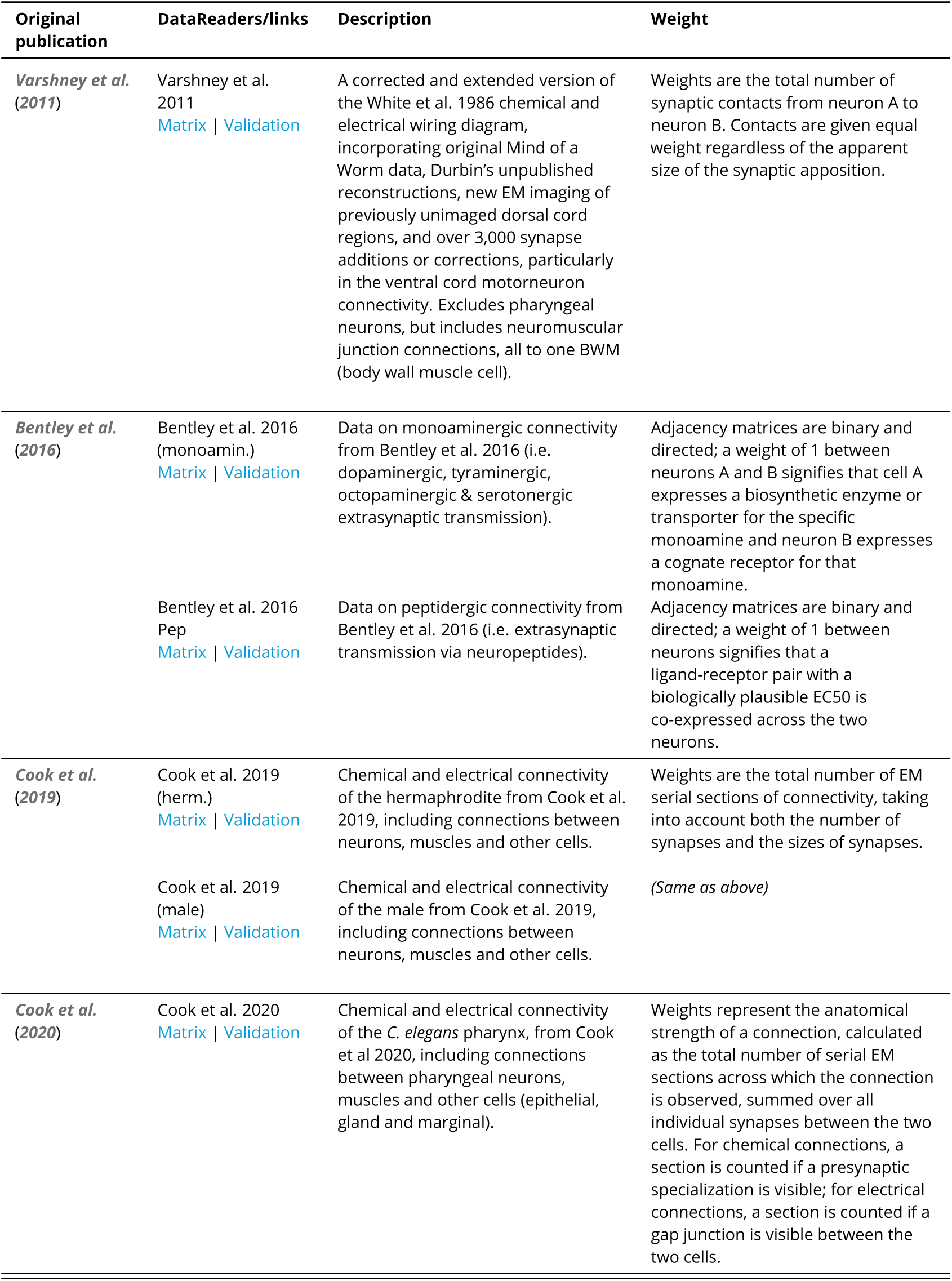
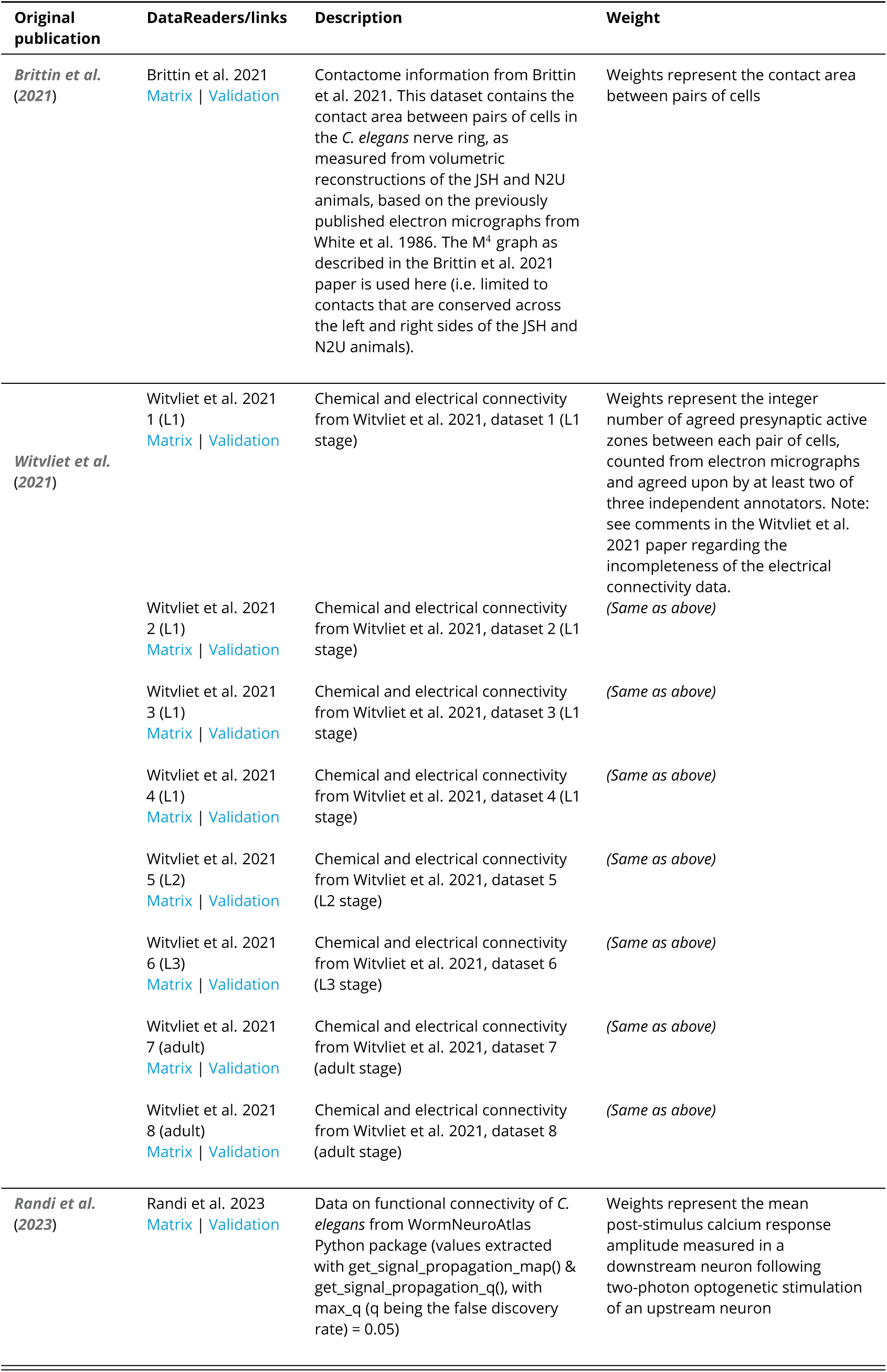
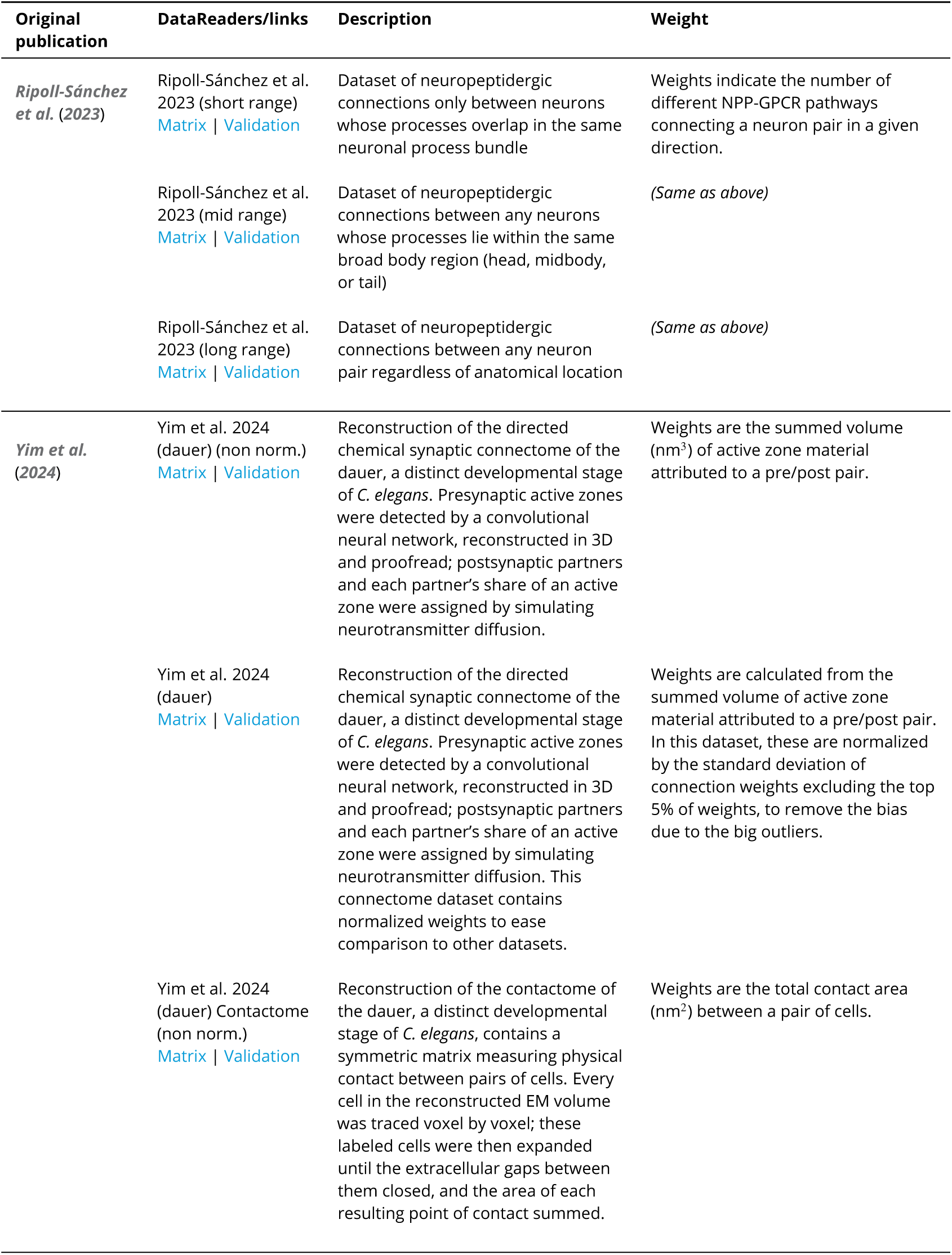
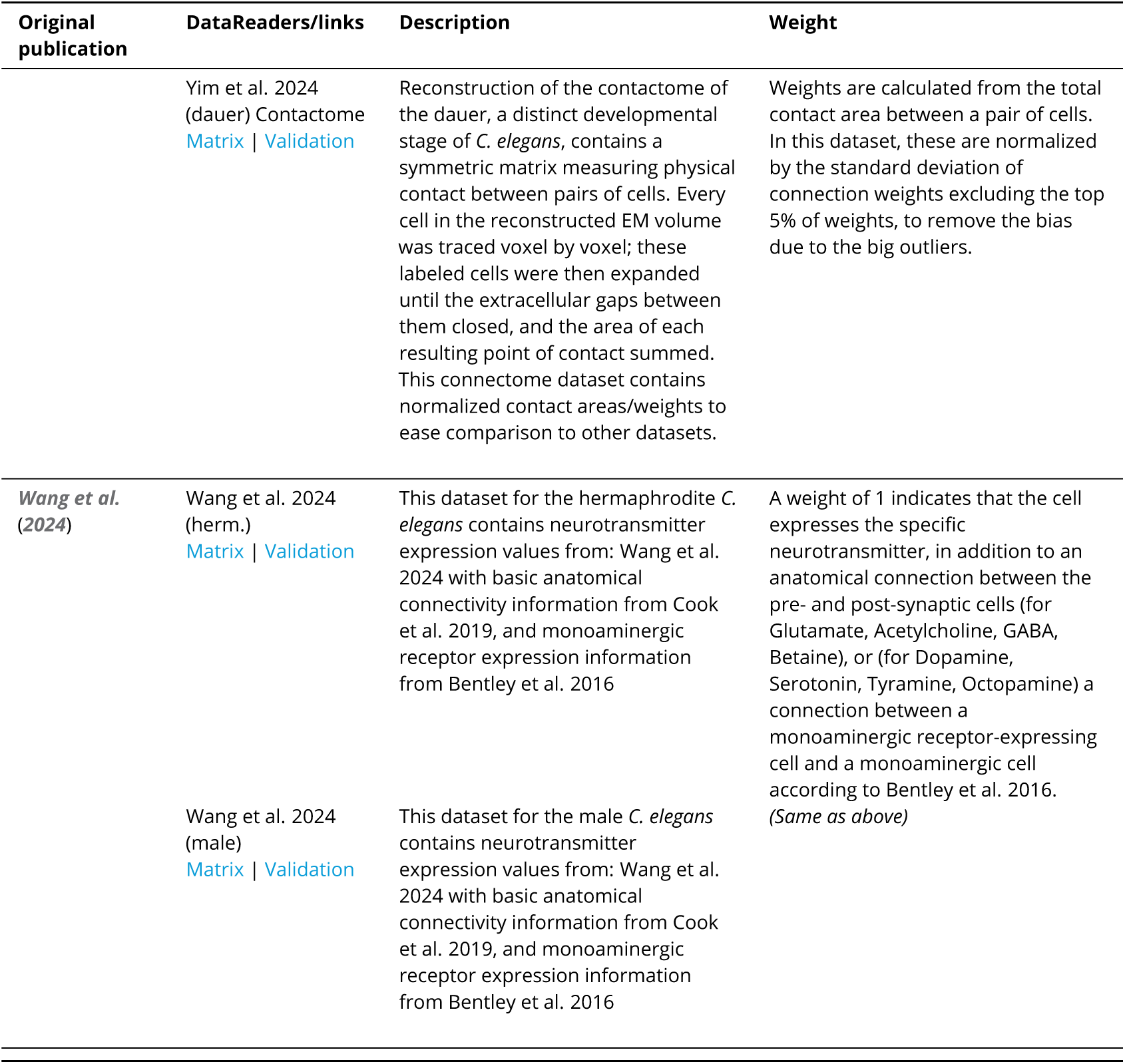
List of all datasets in the *C. elegans* Connectome Toolbox, with descriptions of their DataReaders and interpretations of connectivity weights.

| **Original publication** | **DataReaders/links** | **Description** | **Weight** |
|---|---|---|---|
| ***White et al.*** (***1986***) | White et al. 1986 JSH/L4<br>Matrix \| Validation | Chemical and electrical connectivity of the JSH L4 worm from White et al. 1986 data, based on the neurodata.txt file from R. M. Durbin’s thesis, 1987. See validation information for minor changes from that file. | Weights represent the count of individual presynaptic active zones observed between two neurons across all the serial EM micrographs. |
|  | White et al. 1986 N2U/adult<br>Matrix \| Validation | Chemical and electrical connectivity of the N2U (adult hermaphrodite) worm from White et al. 1986 data, based on the neurodata.txt file from R. M. Durbin’s thesis, 1987. See validation information for minor changes from that file. | *(Same as above)* |
|  | White et al. 1986 (whole worm)<br>Matrix \| Validation | Reanalysis of the White et al. 1986 connectome data by Varshney et al. 2011. This version includes the pharynx. | Used Varshney et al. 2011 interpretation of weights: the total number of synaptic contacts from neuron A to neuron B. Contacts are given equal weight regardless of the apparent size of the synaptic apposition. |
| **Original publication** | **DataReaders/links** | **Description** | **Weight** |
| ***Varshney et al.*** (***2011***) | Varshney et al. 2011<br>Matrix \| Validation | A corrected and extended version of the White et al. 1986 chemical and electrical wiring diagram, incorporating original Mind of a Worm data, Durbin’s unpublished reconstructions, new EM imaging of previously unimaged dorsal cord regions, and over 3,000 synapse additions or corrections, particularly in the ventral cord motorneuron connectivity. Excludes pharyngeal neurons, but includes neuromuscular junction connections, all to one BWM (body wall muscle cell). | Weights are the total number of synaptic contacts from neuron A to neuron B. Contacts are given equal weight regardless of the apparent size of the synaptic apposition. |
| ***Bentley et al.*** (***2016***) | Bentley et al. 2016 (monoamin.)<br>Matrix \| Validation | Data on monoaminergic connectivity from Bentley et al. 2016 (i.e. dopaminergic, tyraminergic, octopaminergic & serotonergic extrasynaptic transmission). | Adjacency matrices are binary and directed; a weight of 1 between neurons A and B signifies that cell A expresses a biosynthetic enzyme or transporter for the specific monoamine and neuron B expresses a cognate receptor for that monoamine. |
|  | Bentley et al. 2016 Pep<br>Matrix \| Validation | Data on peptidergic connectivity from Bentley et al. 2016 (i.e. extrasynaptic transmission via neuropeptides). | Adjacency matrices are binary and directed; a weight of 1 between neurons signifies that a ligand-receptor pair with a biologically plausible EC50 is co-expressed across the two neurons. |
| ***Cook et al.*** (***2019***) | Cook et al. 2019 (herm.)<br>Matrix \| Validation | Chemical and electrical connectivity of the hermaphrodite from Cook et al. 2019, including connections between neurons, muscles and other cells. | Weights are the total number of EM serial sections of connectivity, taking into account both the number of synapses and the sizes of synapses. |
|  | Cook et al. 2019 (male)<br>Matrix \| Validation | Chemical and electrical connectivity of the male from Cook et al. 2019, including connections between neurons, muscles and other cells. | *(Same as above)* |
| ***Cook et al.*** (***2020***) | Cook et al. 2020<br>Matrix \| Validation | Chemical and electrical connectivity of the *C. elegans* pharynx, from Cook et al 2020, including connections between pharyngeal neurons, muscles and other cells (epithelial, gland and marginal). | Weights represent the anatomical strength of a connection, calculated as the total number of serial EM sections across which the connection is observed, summed over all individual synapses between the two cells. For chemical connections, a section is counted if a presynaptic specialization is visible; for electrical connections, a section is counted if a gap junction is visible between the two cells. |
| **Original publication** | **DataReaders/links** | **Description** | **Weight** |
| ***Brittin et al.*** (***2021***) | Brittin et al. 2021<br>Matrix \| Validation | Contactome information from Brittin et al. 2021. This dataset contains the contact area between pairs of cells in the *C. elegans* nerve ring, as measured from volumetric reconstructions of the JSH and N2U animals, based on the previously published electron micrographs from White et al. 1986. The M<sup>4</sup> graph as described in the Brittin et al. 2021 paper is used here (i.e. limited to contacts that are conserved across the left and right sides of the JSH and N2U animals). | Weights represent the contact area between pairs of cells |
| ***Witvliet et al.*** (***2021***) | Witvliet et al. 2021 1 (L1)<br>Matrix \| Validation | Chemical and electrical connectivity from Witvliet et al. 2021, dataset 1 (L1 stage) | Weights represent the integer number of agreed presynaptic active zones between each pair of cells, counted from electron micrographs and agreed upon by at least two of three independent annotators. Note: see comments in the Witvliet et al. 2021 paper regarding the incompleteness of the electrical connectivity data. |
|  | Witvliet et al. 2021 2 (L1)<br>Matrix \| Validation | Chemical and electrical connectivity from Witvliet et al. 2021, dataset 2 (L1 stage) | *(Same as above)* |
|  | Witvliet et al. 2021 3 (L1)<br>Matrix \| Validation | Chemical and electrical connectivity from Witvliet et al. 2021, dataset 3 (L1 stage) | *(Same as above)* |
|  | Witvliet et al. 2021 4 (L1)<br>Matrix \| Validation | Chemical and electrical connectivity from Witvliet et al. 2021, dataset 4 (L1 stage) | *(Same as above)* |
|  | Witvliet et al. 2021 5 (L2)<br>Matrix \| Validation | Chemical and electrical connectivity from Witvliet et al. 2021, dataset 5 (L2 stage) | *(Same as above)* |
|  | Witvliet et al. 2021 6 (L3)<br>Matrix \| Validation | Chemical and electrical connectivity from Witvliet et al. 2021, dataset 6 (L3 stage) | *(Same as above)* |
|  | Witvliet et al. 2021 7 (adult)<br>Matrix \| Validation | Chemical and electrical connectivity from Witvliet et al. 2021, dataset 7 (adult stage) | *(Same as above)* |
|  | Witvliet et al. 2021 8 (adult)<br>Matrix \| Validation | Chemical and electrical connectivity from Witvliet et al. 2021, dataset 8 (adult stage) | *(Same as above)* |
| ***Randi et al.*** (***2023***) | Randi et al. 2023<br>Matrix \| Validation | Data on functional connectivity of *C. elegans* from WormNeuroAtlas Python package (values extracted with get_signal_propagation_map() & get_signal_propagation_q(), with max_q (q being the false discovery rate) = 0.05) | Weights represent the mean post-stimulus calcium response amplitude measured in a downstream neuron following two-photon optogenetic stimulation of an upstream neuron |
| **Original publication** | **DataReaders/links** | **Description** | **Weight** |
| ***Ripoll-Sánchez et al.*** (***2023***) | Ripoll-Sánchez et al. 2023 (short range)<br>Matrix \| Validation | Dataset of neuropeptidergic connections only between neurons whose processes overlap in the same neuronal process bundle | Weights indicate the number of different NPP-GPCR pathways connecting a neuron pair in a given direction. |
|  | Ripoll-Sánchez et al. 2023 (mid range)<br>Matrix \| Validation | Dataset of neuropeptidergic connections between any neurons whose processes lie within the same broad body region (head, midbody, or tail) | *(Same as above)* |
|  | Ripoll-Sánchez et al. 2023 (long range)<br>Matrix \| Validation | Dataset of neuropeptidergic connections between any neuron pair regardless of anatomical location | *(Same as above)* |
| ***Yim et al.*** (***2024***) | Yim et al. 2024 (dauer) (non norm.)<br>Matrix \| Validation | Reconstruction of the directed chemical synaptic connectome of the dauer, a distinct developmental stage of *C. elegans*. Presynaptic active zones were detected by a convolutional neural network, reconstructed in 3D and proofread; postsynaptic partners and each partner’s share of an active zone were assigned by simulating neurotransmitter diffusion. | Weights are the summed volume (nm<sup>3</sup>) of active zone material attributed to a pre/post pair. |
|  | Yim et al. 2024 (dauer)<br>Matrix \| Validation | Reconstruction of the directed chemical synaptic connectome of the dauer, a distinct developmental stage of *C. elegans*. Presynaptic active zones were detected by a convolutional neural network, reconstructed in 3D and proofread; postsynaptic partners and each partner’s share of an active zone were assigned by simulating neurotransmitter diffusion. This connectome dataset contains normalized weights to ease comparison to other datasets. | Weights are calculated from the summed volume of active zone material attributed to a pre/post pair. In this dataset, these are normalized by the standard deviation of connection weights excluding the top 5% of weights, to remove the bias due to the big outliers. |
|  | Yim et al. 2024 (dauer) Contactome (non norm.)<br>Matrix \| Validation | Reconstruction of the contactome of the dauer, a distinct developmental stage of *C. elegans*, contains a symmetric matrix measuring physical contact between pairs of cells. Every cell in the reconstructed EM volume was traced voxel by voxel; these labeled cells were then expanded until the extracellular gaps between them closed, and the area of each resulting point of contact summed. | Weights are the total contact area (nm<sup>2</sup>) between a pair of cells. |
| **Original publication** | **DataReaders/links** | **Description** | **Weight** |
|  | Yim et al. 2024 (dauer) Contactome<br>Matrix \| Validation | Reconstruction of the contactome of the dauer, a distinct developmental stage of *C. elegans*, contains a symmetric matrix measuring physical contact between pairs of cells. Every cell in the reconstructed EM volume was traced voxel by voxel; these labeled cells were then expanded until the extracellular gaps between them closed, and the area of each resulting point of contact summed. This connectome dataset contains normalized contact areas/weights to ease comparison to other datasets. | Weights are calculated from the total contact area between a pair of cells. In this dataset, these are normalized by the standard deviation of connection weights excluding the top 5% of weights, to remove the bias due to the big outliers. |
| ***Wang et al.*** (***2024***) | Wang et al. 2024 (herm.)<br>Matrix \| Validation | This dataset for the hermaphrodite *C. elegans* contains neurotransmitter expression values from: Wang et al. 2024 with basic anatomical connectivity information from Cook et al. 2019, and monoaminergic receptor expression information from Bentley et al. 2016 | A weight of 1 indicates that the cell expresses the specific neurotransmitter, in addition to an anatomical connection between the pre- and post-synaptic cells (for Glutamate, Acetylcholine, GABA, Betaine), or (for Dopamine, Serotonin, Tyramine, Octopamine) a connection between a monoaminergic receptor-expressing cell and a monoaminergic cell according to Bentley et al. 2016. |
|  | Wang et al. 2024 (male)<br>Matrix \| Validation | This dataset for the male *C. elegans* contains neurotransmitter expression values from: Wang et al. 2024 with basic anatomical connectivity information from Cook et al. 2019, and monoaminergic receptor expression information from Bentley et al. 2016 | *(Same as above)* |

To support accessible interactive exploration of connectomic data by users, we developed general Python visualization and analysis methods that work with any ConnectomeDatasets. Since these datasets can contain hundreds of cells and thousands of connections, we defined a number of ConnectomeView classes, which describe subsets of cells (or lists of nodes containing grouped cells), e.g. just the pharyngeal neurons, or grouped interneuron/motorneuron/sensory neuron views. This allows the same aspects of different datasets to be visualized in a common way. Each view can in turn be represented in a number of graphical formats. Fig. 3, supplement 1 shows a number of these for a single dataset (***Varshney et al., 2011***), including: adjacency matrices visualized in 2D heatmaps, where cells are grouped by type (e.g. sensory, interneuron, motorneuron) and the color shows the relative weight of the connection; force-directed graphs showing the interconnected cells; circuit views illustrating the connectivity between grouped cells; and hive plots, showing connectivity between and within the 3 main subtypes of neurons. Each of these graphical types (described in more detail below) can be generated for each of the ConnectomeViews of the individual datasets (as long as they are within scope of the view, e.g. the pharynx view will not be appropriate for datasets missing pharyngeal neurons).

Additionally, while these graphs can be generated by researchers locally using the Python framework (e.g. in a Jupyter notebook, Fig. 2, supplement 3), they have also been pre-generated in each combination for quick access on the website (Fig. 2, supplement 1, https://openworm.org/ConnectomeToolbox). The website also summarizes all data related to each neuron, muscle or other connected cell (Fig. 2, supplement 2).

### Connectome Toolbox consolidates anatomical connectomic data

A central aim of the Connectome Toolbox is to consolidate fragmented anatomical connectome datasets into a unified, machine-readable, easily extensible framework that enables direct comparison across studies. Multiple anatomical reconstructions of the *C. elegans* nervous system have been published over the past four decades, differing in biological scope, reconstruction methodology, and annotation conventions (***White et al., 1986***; ***Varshney et al., 2011***; ***Cook et al., 2019***, ***2020***; ***Witvliet et al., 2021***; ***Yim et al., 2024***). These datasets vary in the number of neurons included, the types of connections represented (chemical synapses, gap junctions, or non-neuronal targets), and the developmental or sex-specific context in which they were generated. As a result, direct comparison across studies is non-trivial, and apparent conflicts between connectomes are often caused by differences in scope of the studies or the presentation of the data, rather than genuine biological inconsistencies.

To address this fragmentation, the Connectome Toolbox consolidates anatomical connectomic data from multiple published sources within a common reference-aligned framework. We adopt the neuron naming and annotation conventions of ***Cook et al. (2019)*** as a unifying scaffold, as this reconstruction currently provides the most comprehensive coverage of neuron classes, connection types, and synaptic targets available (***Cook et al., 2019***). Importantly, this choice does not treat any single dataset as definitive; rather, it provides a practical reference against which smaller, earlier, or modality-specific datasets can be mapped and compared while preserving their original provenance. Comparison of anatomical connectivity across datasets reveals a clear subset structure, with ***Cook et al. (2019)*** representing the upper bound of reported connections. As shown in Fig. 3, supplement 2, earlier anatomical reconstructions consistently contain fewer connections per neuron class, while the ***Cook et al. (2019)*** dataset encompasses almost all connections present in these datasets and includes additional synapses and targets absent from more limited reconstructions. The adult male connectome from ***Cook et al. (2019)*** forms a distinct case, reflecting genuine biological differences rather than reconstruction scope. Note that datasets can be compared directly in terms of which connections are present between pairs of cells, but not in terms of edge weights, as each publication defines weight differently (e.g. active zone counts, serial sections, contact area). See Table 1 for the specific interpretation of weight used in each dataset. This consolidated anatomical representation underpins the subsequent analyses in which anatomical connectivity is integrated with molecular annotations (e.g. neurotransmitter expression), functional data, and computational models.

### Connectome Toolbox provides curated data validated against published connectomes

An important goal of the Connectome Toolbox is to provide a reliable, reproducible, and validated representation of published connectome datasets. This will increase the FAIRness of the datasets, i.e. ensuring they are Findable, Accessible, Interoperable, and Reusable for other scientific investigations (***Wilkinson et al., 2016***). ***Supplementary file 1*** describes in detail the curation process for each dataset we have incorporated, with links to the original data sources (e.g. the supplementary information files of the original publication, or obtained from sites such as https://wormwiring.org), cataloging any inconsistencies/discrepancies found in the original data files and how they were addressed, and linking to the final files used in our DataReader classes.

Additionally, a set of machine-readable validation tests have been generated for each dataset, which allow automated verification that the imported data in the ConnectomeDataset, and hence accessed through the Connectome Toolbox API, is consistent with the original published data. Checks here can include total connection numbers, weights of specific connections and total connection weights (which are extracted from the original data sources, e.g. Excel spreadsheets), as well as consistency checks, such as ensuring each electrical connection between two cells is represented in both directions, and the weights should be equal. These validation tests are run every time the web interface is updated, and the results are logged and made available to users.

### Connectome Toolbox generates graphical views to facilitate exploration and interpretation of connectomics data

Traditionally, connectomes are often represented as either matrices or connected graphs (whose nodes can represent individual cells, cell classes or larger groups of cells in anatomical brain regions). We outline here how various graphical views of datasets we have assembled, including hive plots as an alternative approach to visualizing connectomics data, can be generated using the toolbox.

### Adjacency matrices

All of the connectivity datasets can be represented as adjacency matrices, where the rows represent presynaptic cells and the columns represent postsynaptic cells. The value in each cell of the matrix represents the weight of the connection between the cells, which has different interpretations depending on the dataset, e.g. numbers of synapses identified between pairs of cells in the anatomical connectomes, or strength of connections (positive or negative) in functional connectomes (Table 1). Fig. 4a shows the chemical connections in the hermaphrodite connectome from ***Cook et al. (2019)***. Cells are grouped by type, alphabetically within groups, and colored according to the type of cell (based on usage in WormAtlas (***Altun et al., 2025***), see Appendix 1 Fig. 1). The presence of multiple non-neuronal cells (many of which are muscles, shown in green) distinguishes this view from the matrices shown in Fig. 3, supplement 1a,b. Only a subset of cells in this dataset have been labeled on the axes for clarity. High level connectivity patterns can be observed from this view, including dense connections from both sensory neurons (pink) and interneurons (red) to interneurons, with both groups also projecting to sensory and motorneurons (purple). Motorneurons mainly target themselves and body wall muscles (dark green).

**Figure 4.**
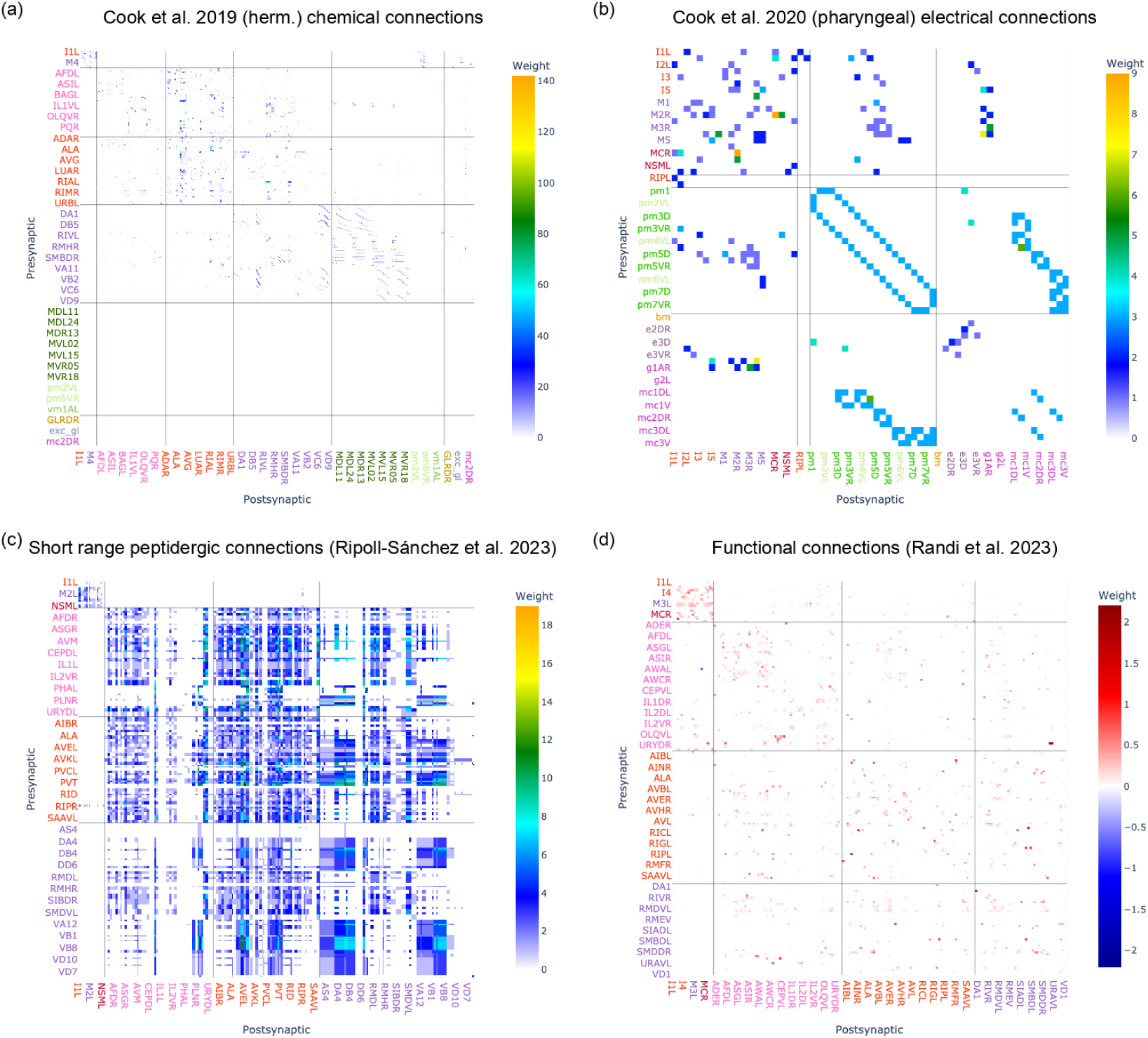
Visualizing connectivity in datasets with adjacency matrices. (a) All chemical connections in the hermaphrodite connectome of ***Cook et al. (2019)***. (b) All electrical connections in the hermaphrodite pharynx from ***Cook et al. (2020)***. (c) Short range peptidergic connections from ***Ripoll-Sánchez et al. (2023)***. (d) Functional connections from ***Randi et al. (2023)***, showing which cells are activated (red) or inhibited (blue) following optogenetic stimulation of the presynaptic cell. Colors of cell names in the axes labels specify their types (e.g. sensory (pink), interneuron (red), motorneuron (purple), muscle (green)), which are outlined in full in Appendix 1 Fig. 1. Not all cell names are shown on axes for clarity. In the online interactive versions of these figures, hovering over the square representing any of the connections will show the pre and postsynaptic cell names and weight of the connection, and zooming in will reveal more cell names on axes. More details on each of the datasets, as well as their individual definitions of “weight”, can be found in Table 1.

Fig. 4b shows the electrical connections in the hermaphrodite pharynx from ***Cook et al. (2020)***. Different connectivity patterns are evident between the neuronal cells (top left) and the surrounding pharyngeal muscle cells (pm-), and support cells (epithelial, e-, gland, g- and marginal cells, mc-). Fig. 4c shows the short range peptidergic connections from ***Ripoll-Sánchez et al. (2023)***. Here, the matrix view is particularly useful as the dense connectivity makes graph views difficult to interpret. Fig. 4d depicts functional connections from ***Randi et al. (2023)***, with connection weights showing whether optogenetic activation of a presynaptic cell results in activation (red) or inhibition (blue) of the postsynaptic cell. This is one example of a connectivity matrix from this dataset, with maximum q value (false discovery rate, configurable in the DataReader class) set to 0.05 (Table 1). In this way, multiple connectivity matrices can be generated from the same dataset, and corresponding adjacency matrix visualizations created, allowing users to explore how the connectivity changes with different thresholds for significance.

Fig. 2, supplement 1 shows an adjacency matrix view of the dauer contactome from ***Yim et al. (2024)***, which is symmetric, and contains a much larger number of contacts (not all of them synaptic) between the cells.

### Graph based views

Graph-based visualization provides an intuitive and powerful way to explore neuronal connectivity within the *C. elegans* nervous system. Graphs represent neurons, muscles or other connected cells as nodes, while the connections between them (chemical synapses, gap junctions, or other forms of interaction) are represented as edges. This representation not only mirrors the biological structure of the nervous system but also provides a foundation for applying graph-theoretic methods to analyze network topology, modularity, and information flow (***Emmons, 2024***).

Within the Connectome Toolbox, graph visualizations are implemented using the NetworkX package (***Hagberg et al., 2008***) and can be generated for any dataset or defined subset of cells. The use of a force-directed layout allows clusters of highly connected neurons to emerge naturally, revealing the organizational principles of the connectome. This view is particularly valuable for identifying subnetworks, functional motifs, and inter-system linkages that may be less apparent in matrix-based representations.

As shown in Fig. 5, illustrating chemical and electrical synaptic connectivity from ***Cook et al. (2019)***, the *C. elegans* nervous system is composed of two major domains: the large somatic nervous system and the smaller, relatively autonomous pharyngeal nervous system. The graph view clearly illustrates these as distinct but interconnected subnetworks, linked by a small number of “bridge” neurons that mediate communication between them (the interactive view of this graph can be used to easily identify these as connections to RIPL/RIPR). This view can be contrasted with the ***Varshney et al. (2011)*** data in Fig. 3, supplement 1c, where the pharyngeal neuron connectivity is not included, and the pharyngeal neurons form an isolated cluster on the left of the graph. The inclusion of muscle cells in the ***Cook et al. (2019)*** dataset extends this framework beyond neurons, highlighting the integration of neuronal and effector networks, e.g. gap junctions between adjacent body wall muscles (dark green in Fig. 5b,d) and to and between vulval/uterine muscles (light green in Fig. 5b).

**Figure 5.**
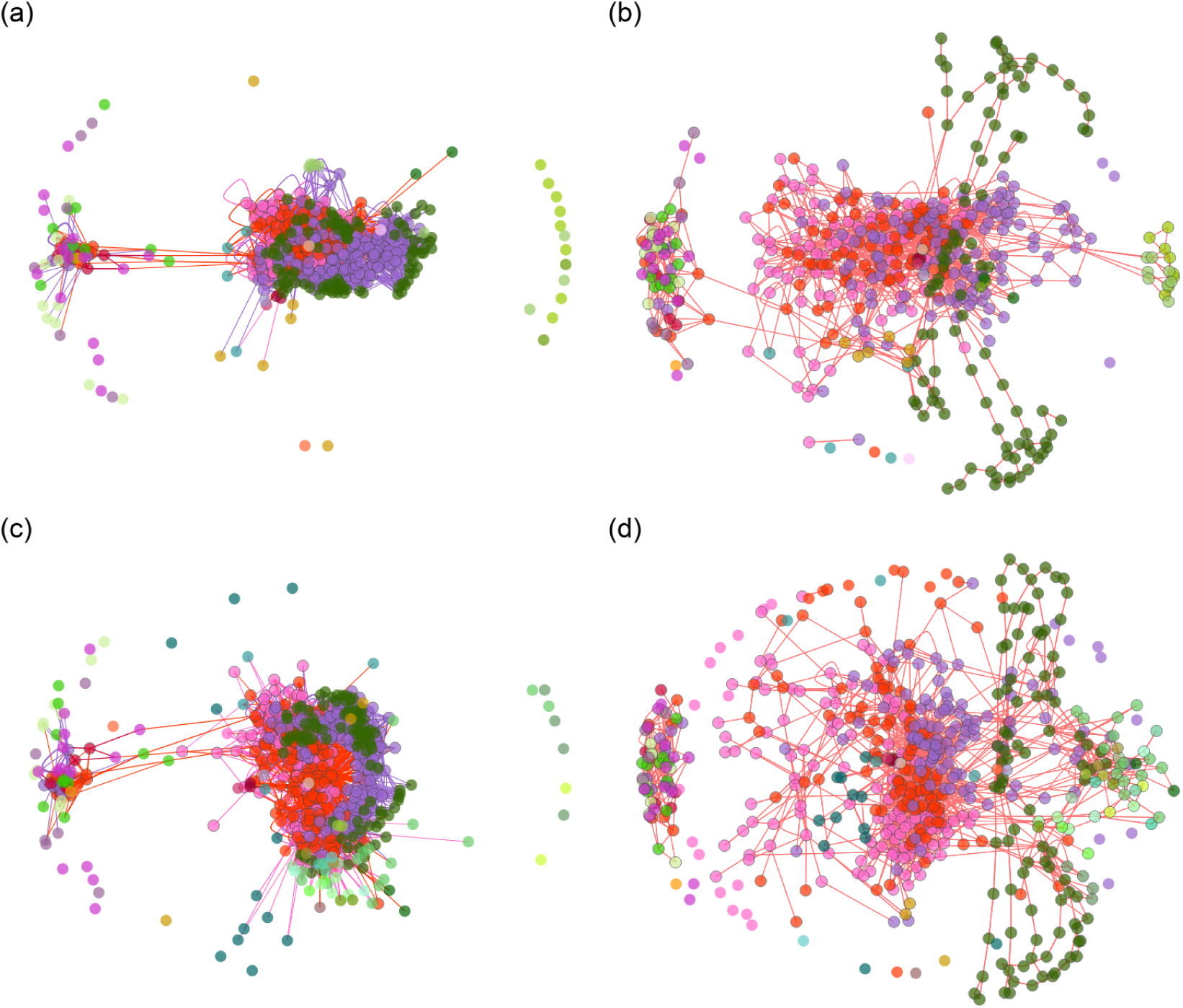
Force-directed graphs of network connectivity from ***Cook et al. (2019)***. (a) Hermaphrodite chemical connections. (b) Hermaphrodite electrical connections. (c) Male chemical connections. (d) Male electrical connections. Body wall muscles are in dark green, other muscles in light green, e.g. vulval/uterine muscles on right in (a) and (b). Colors of nodes are given in Appendix 1 Fig. 1.

In addition to providing a structural overview of connectivity, the graph view enables functional interpretations of information flow. The directional organization of the network—from sensory (pink) to interneuron (red) to motorneuron (purple) and muscle (green) connectivity—can be readily observed, reflecting the canonical flow of neural processing underlying behavior.

Graph based visualizations also provide specific small circuit views where multiple cells are grouped into nodes and the aggregated connections between them are represented as edges. This approach allows for a more focused analysis of specific physiologically relevant circuits, such as the escape response circuit (***Pirri and Alkema, 2012***) shown in Fig. 3, supplement 1d, or more compact high level views of the connectivity across the whole network (Fig. 3, supplement 1e).

### Hive plots

While traditional force-directed graph layouts are useful for intuitively visualizing connectivity/high-level organization and identifying clusters, they suffer from a key limitation: their spatial arrangement is non-deterministic. The same network can appear visually different each time it is plotted, making direct comparison across datasets or between network subsets difficult. Hive plots (***Krzy-winski et al., 2012***) provide a complementary, rule-based approach to network visualization that addresses this limitation by assigning nodes to fixed axes according to defined biological or topological properties. Fig. 6, supplement 1 outlines the core concepts of hive plots.

**Figure 6.**
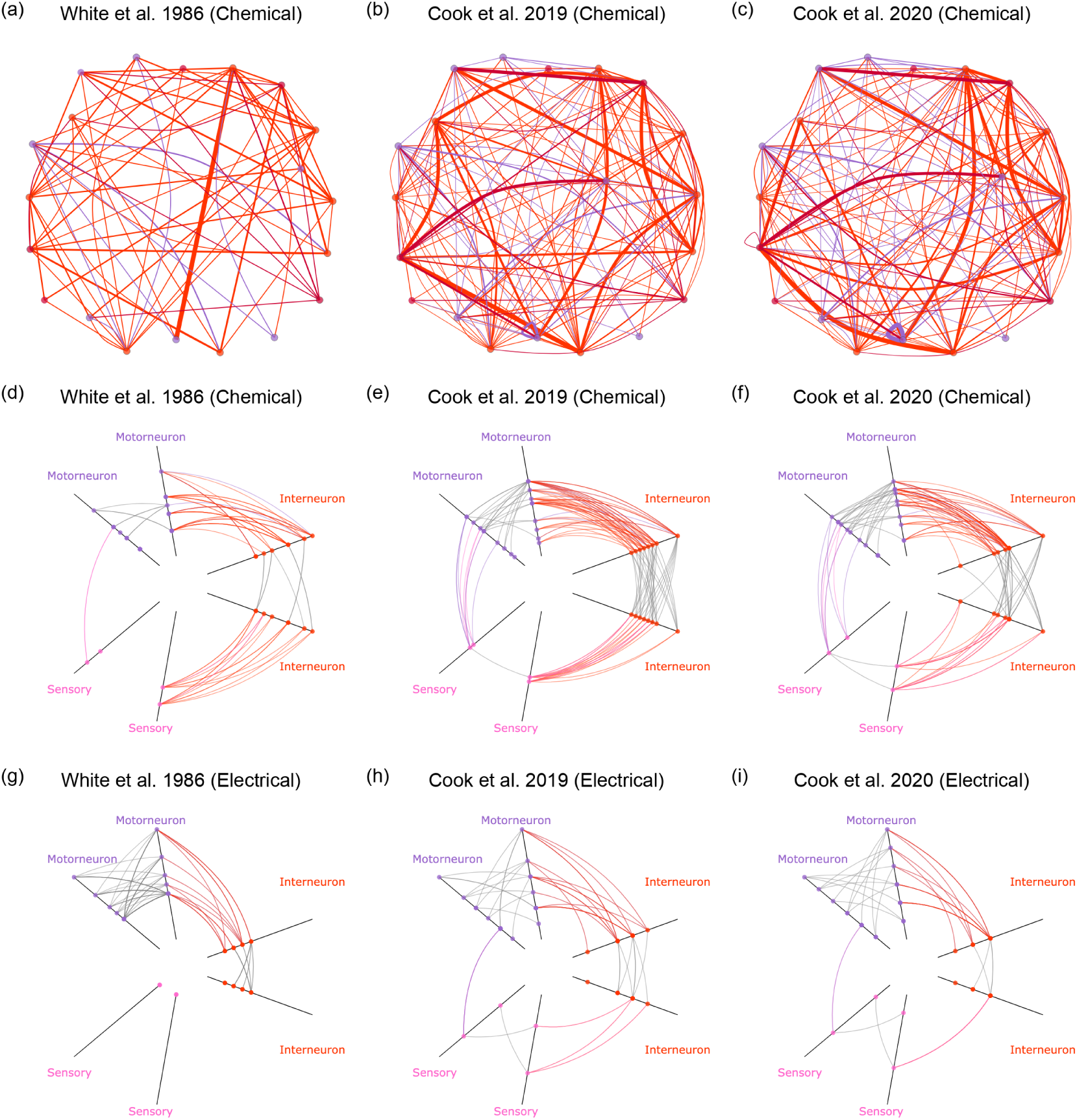
Using hive plots to compare synaptic connectivity in the pharynx across multiple connectome datasets. (a-c) Force-directed graph views of the chemical connections between sensory neurons (pink), interneurons (red) and motorneurons (purple) in the pharyngeal nervous system from three different datasets: (a) ***White et al. (1986)*** (72 individual connections between neurons), (b) ***Cook et al. (2019)*** (141 connections) and (c) ***Cook et al. (2020)*** (148 connections). (d-f) Corresponding hive plots of chemical connections from the same three datasets. (g-i) Corresponding hive plots of electrical connections from the same three datasets (92, 66 and 54 electrical connections respectively). All panels are screenshots of interactive plots from the website, where hovering over the nodes reveals their cellular identities, as well as provides information about the numbers of incoming and outgoing connections. **Figure 6–Figure supplement 1**. Overview of hive plots.

In a hive plot, each axis represents a distinct category of the nodes, such as sensory, interneuron, and motorneuron classes, and nodes themselves are positioned along each axis based on quantitative attributes such as degree, connection strength, or anatomical position. Edges between nodes are then drawn as curves linking the axes, resulting in a reproducible and interpretable representation of network structure. This deterministic layout facilitates direct comparison between datasets, enabling researchers to visualize how patterns of connectivity differ between sexes, developmental stages, or connectivity modalities (e.g., anatomical versus functional). Hive plots are particularly effective for highlighting global organizational principles such as feedforward versus recurrent wiring, cross-modality interactions, or long-range connections that bridge neuronal classes.

Previous work has demonstrated the power of hive plots for revealing non-canonical pathways in *C. elegans*. For example, Bentley and colleagues (***Bentley et al., 2016***) used this approach to show how monoaminergic signaling can bypass premotor interneurons and directly modulate motorneurons, revealing parallel routes of sensory information flow. Fig. 6 illustrates examples of hive plots we have generated from several datasets containing connectivity of the pharyngeal nervous system (***White et al., 1986***; ***Cook et al., 2019***, ***2020***). These visualizations show a clearer depiction of the high level connectivity between different classes of cells in the pharynx compared to the “hairball” of the force-directed graphs (Fig. 6a-c), and how the distinct patterns of both chemical (Fig. 6d-f) and electrical connectivity (Fig. 6g-i) are preserved across each dataset. An example of a change across these datasets which can easily be determined from these views is that the later two datasets (Fig. 6e,f) show greater chemical connectivity between neurons classed as sensory and motorneurons compared to more standard feedforward sensory-interneuron-motorneuron pathway of the original ***White et al. (1986)*** dataset (Fig. 6d). This change, however, is not reflected in the electrical connectivity, which remains primarily either between motorneurons or between motorneurons and interneurons (Fig. 6g-i).

Fig. 3, supplement 1f shows another example of a hive plot, this time for the full chemical synaptic connectome of the somatic nervous system described in ***Varshney et al. (2011)***. This highlights how highly connected “rich club” neurons (***Towlson et al., 2013***) such as AVAL/R can easily be identified from this view.

These examples underscore the value of Connectome Toolbox hive plots as a reproducible, quantitative, and comparative framework for exploring complex neuronal networks.

### Connectome Toolbox provides a unified programmatic interface integrating anatomical and neurotransmitter data

We have combined the anatomical wiring data from ***Cook et al. (2019)*** and a map of monoaminergic connections from ***Bentley et al. (2016)***, with the recently published comprehensive neuro-transmitter atlas (***Wang et al., 2024***), enabling direct exploration of how transmitter identity is distributed across the wiring diagram, and how molecular signaling systems map onto circuit architecture in *C. elegans*. ***Wang et al. (2024)*** presented a comprehensive, CRISPR-based fluorescent reporter atlas mapping all major neurotransmitter systems (the biosynthetic and vesicular machinery) across both male and hermaphrodite *C. elegans*, revealing new neurotransmitter expression sites, co-transmission patterns, and sex-specific signaling. The integration of the neurotransmitter identities onto the anatomical wiring data (***Cook et al., 2019***) allows the anatomical reach of each transmitter system to be traced through the connectome, i.e. which cells release it, which cells they contact, and where different systems converge on common targets. Additionally, we have integrated the monoaminergic connectome from ***Bentley et al. (2016)*** which provides a map of extrasynaptic connections mediated by serotonin, dopamine, tyramine, and octopamine, using the updated ***Wang et al. (2024)*** data for the neurotransmitter identities of the presynaptic neurons.

Displaying these systems across the full connectome is impractical: with several hundred cells and several thousand connections, the resulting graphs are dense enough that the pattern specific to any one transmitter is lost among the others. Connectome Toolbox addresses this through ConnectomeView classes, which aggregate cells into a smaller number of named nodes and sum the connections between them, so that a whole-network pattern can be read at a glance while the underlying cell-level data remain unchanged. To illustrate how this can be used to explore the integrated dataset, we present a series of visualizations that overlay neurotransmitter systems onto two such higher-level circuit views. Fig. 7 uses a layout and grouping of neurons based on Fig. 3 of ***Cook et al. (2019)***; Fig. 7, supplement 2 uses the nerve-ring map of ***Brittin et al. (2021)***, known as “Brainmap”.

**Figure 7.**
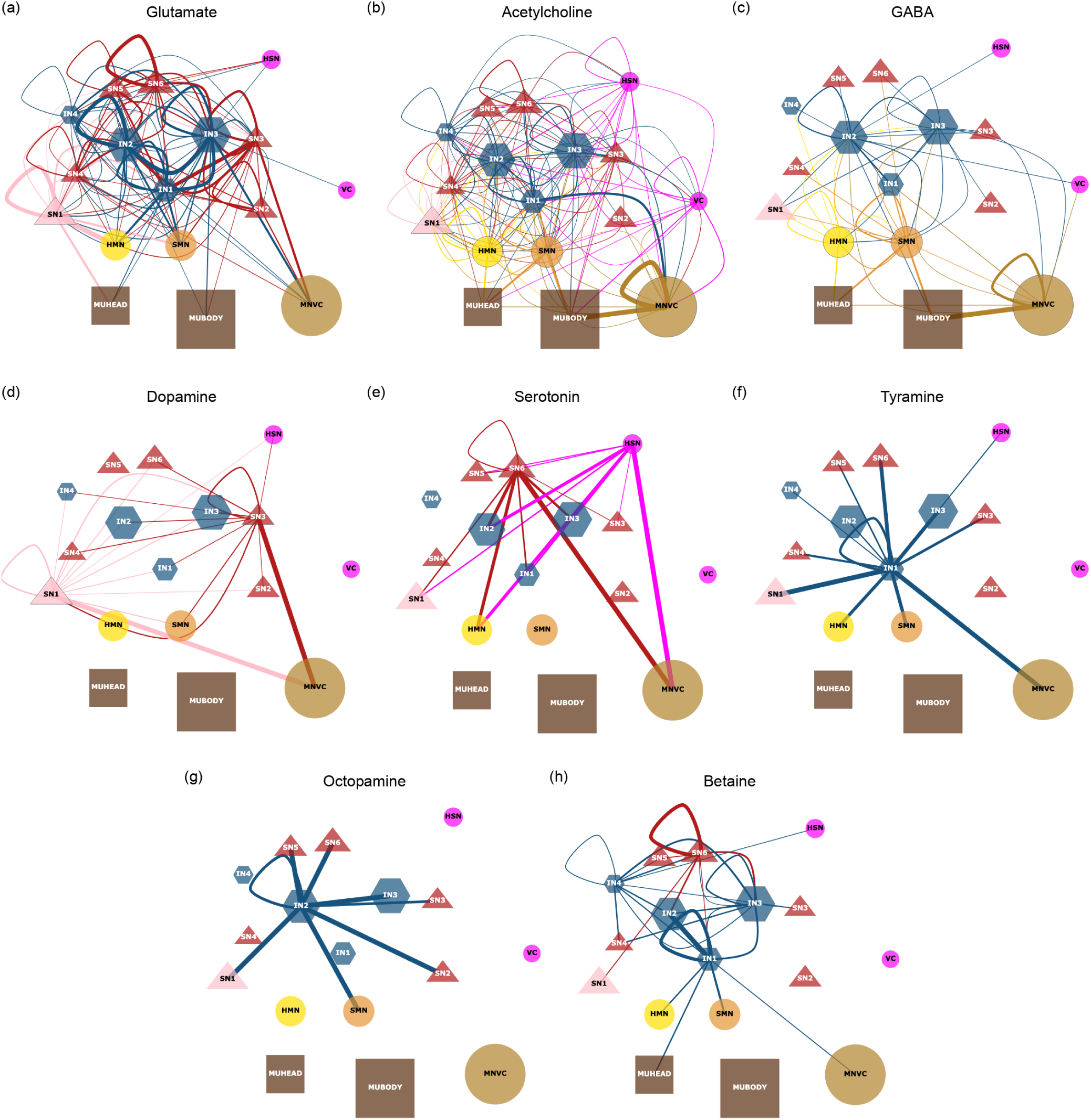
Visualizations of neurotransmitter sources from *Wang et al. (2024)* integrated with the anatomical connectome from *Cook et al.* (*2019*) and the monoaminergic connectome from *Bentley et al.* (*2016*). The full ConnectomeDataset contains 8 connectivity matrices, one for each neurotransmitter, with the presynaptic neurons defined by the neurotransmitter expression from *Wang et al. (2024)* and the postsynaptic cells (both neurons and muscles) defined by the presence of a chemical synaptic connection in *Cook et al.* (*2019*) (for glutamate, acetylcholine, GABA and betaine) or by the presence of a monoaminergic connection in *Bentley et al.* (*2016*) (for serotonin, dopamine, tyramine and octopamine). **(a-h)** ConnectomeView representations of the connectivity for each neurotransmitter using a grouping of cells based on Fig. 3 in *Cook et al.* (*2019*). Figure Supplement 1 describes the groups, lists the cells present in each, and specifies their colors, including: blue - interneurons (IN*); red/pink - sensory neurons (SN*); light brown - motorneurons of the ventral cord (MNVC); dark brown - muscles (MU*). Connecting lines are colored by the group containing the neurons expressing the neurotransmitter, and the thickness of the line is proportional to the total number of connections between pairs of cells in the two groups. Note that the scaling of line width is per panel. **(a)** Glutamate. **(b)** Acetylcholine. **(c)** GABA. **(d)** Dopamine. **(e)** Serotonin. **(f)** Tyramine. **(g)** Octopamine. **(h)** Betaine. **Figure 7–Figure supplement 1.** Screenshot of *Cook et al.* (*2019*) Fig. 3 ConnectomeView webpage, describing groups and listing cells in each. **Figure 7–Figure supplement 2.** Brainmap (*Brittin et al., 2021*) based view of neurotransmitters

Glutamate is a major excitatory neurotransmitter, with the vesicular reporter allele of *eat-4/VGLUT* defining glutamatergic neurons (***Serrano-Saiz et al., 2013***; ***Wang et al., 2024***). It can also activate glutamate-gated chloride channels leading to inhibition of downstream motorneurons or muscles in locomotion (***Horoszok et al., 2001***; ***Sato et al., 2021***). Fig. 7a and Fig. 7, supplement 2a show the broad distribution of glutamatergic neurons across the *C. elegans* connectome. Due to its high number of connections, it is difficult to see the exact pattern of connectivity. However, in Fig. 7a, the strong connectivity between interneuron groups can clearly be observed.

Acetylcholine, identified by a reporter allele for *unc-17/VAChT*, is found to be the most widely used neurotransmitter with excitatory and inhibitory roles (***Pereira et al., 2015***; ***Wang et al., 2024***). Because both major excitatory transmitters can also act through inhibitory receptors, inferring the sign of circuit connections from transmitter identity alone is difficult (***Sato et al., 2021***; ***Kumar et al., 2024***). Fig. 7b and Fig. 7, supplement 2b show the widespread distribution of cholinergic neurons. A significant portion of motorneurons are cholinergic (Fig. 7b, bottom right), consistent with their role in driving muscle contraction (***Pereira et al., 2015***).

GABA (*γ*-aminobutyric acid) is the major inhibitory neurotransmitter and is defined by the expression of either *unc-25/GAD* or *unc-47/VGAT* (***Gendrel et al., 2016***). GABAergic signaling is critical for producing sinusoidal movement of the worms and also involved in egg laying, sensory gating as well as behavioral state transitions. Fig. 7c highlights the relatively limited number of GABAergic connections in the *C. elegans* connectome, again with a large portion of these coming from motorneurons. In Fig. 7, supplement 2, it can be more clearly seen that AVJ, RIB and ventral cord motorneurons act as hubs for GABAergic signaling.

Overlaying monoaminergic systems - dopamine, serotonin, tyramine and octopamine - onto the synaptic wiring reveals distinct modulatory architectures as shown in Fig. 7d-g and Fig. 7, supplement 2d-g. The important work of ***Bentley et al. (2016)*** led to dissecting the connectome into multiple layers, thus demonstrating the alternative modes of interaction between neurons. The transmission of these monoamines is often slower, and modulate activity on non-synaptic sites.

Dopamine, a catecholamine synthesized by *cat-2/TH*, contributes to behavioral plasticity of *C. elegans*. Fig. 7d and Fig. 7, supplement 2d highlight the main dopaminergic neurons in *C. elegans*, projecting from sensory neurons such as CEP and ADE to many populations, particularly motorneurons.

Serotonin is a neuromodulatory monoamine synthesized by *tph-1/TPH* and *bas-1/AAAD*, and it regulates locomotion by causing feeding-induced slowing (***Dag et al., 2023***). Fig. 7e shows groups SN6 and HSN (hermaphrodite-specific neurons) as hubs for serotonergic signaling. Fig. 7, supplement 2e lacks HSN neurons, though the strong serotonergic connections to/from the pharynx can be seen, with the NSM neurons playing a key role in modulating feeding behavior (***Dag et al., 2023***).

Tyramine and octopamine are defined by markers for *tdc-1/TDC* and *tbh-1/TBH*, with tyramine (defined by *tdc-1+/tbh-1-*) having a key role in locomotion and escape responses, whereas octopamine (*tdc-1+/tbh-1+*) modulates arousal, foraging and stress-related behaviors. Fig. 7f and Fig. 7g highlight the relatively limited number of tyraminergic neurons (RIM in layer 1 interneurons - IN1) and octopaminergic neurons (RIC in IN2) in *C. elegans*. The specific neurons are more clearly seen in Fig. 7, supplement 2f and Fig. 7, supplement 2g.

Betaine, also known as trimethylglycine, is a less well studied signaling molecule in *C. elegans*, established as a neuronal transmitter only relatively recently and with its range of action across the nervous system still incompletely mapped. It acts on two distinct, independently evolved ligand-gated ion channels with opposing effects: the betaine-gated chloride channel LGC-41, which mediates inhibitory signaling and is expressed broadly across the navigation circuit (***Hardege et al., 2022***), and the betaine-gated cation channel ACR-23, which is excitatory and is related to nicotinic acetylcholine receptors (***Peden et al., 2013***).

Our integration of the betaine data from ***Wang et al. (2024)*** onto the anatomical connectome (Fig. 7h) shows the anatomical output projections of neurons that co-express the betaine uptake transporter snf-3/BGT1 and the vesicular monoamine transporter cat-1/VMAT (in the hermaphrodite: ASI, AUA, NSM, RIM and RIR). These neurons can therefore take up betaine and potentially release it via vesicles, and were identified by ***Wang et al. (2024)*** as candidate betaine-signaling neurons. Neurons expressing snf-3/BGT1 without cat-1/VMAT were excluded; ***Wang et al. (2024)*** propose that these act in betaine clearance rather than release. We note that this atlas maps the machinery for betaine uptake and release rather than the betaine receptors LGC-41 and ACR-23. The panel therefore identifies the anatomical partners of betaine-associated neurons, not cells demonstrated to be betaine-responsive.

Whereas LGC-41 controls behavioral-state transitions during food search, ACR-23 has been characterized mainly in mechanosensory neurons, where it maintains basal levels of locomotion (***Peden et al., 2013***). Neuronally produced betaine, released from the RIM interneurons and acting via LGC-41, has been identified as an inhibitory neuromodulator critical for the switch from local to global foraging behavior (***Hardege et al., 2022***). Fig. 7h and Fig. 7, supplement 2h place this model in an anatomical context: they show which components of the local-search circuit are postsynaptic partners of RIM, and thus candidate sites at which RIM-derived betaine could act. Because the overlay encodes neither betaine-receptor expression nor connection sign, it delineates the anatomical reach of betaine-associated neurons across the navigation network, defining a set of downstream candidates for betainergic modulation, rather than demonstrating that any given target is inhibited.

This betaine-dependent modulation complements the parallel-circuit framework proposed for foraging-state transitions, in which chemosensory and mechanosensory glutamatergic inputs redundantly suppress the integrating neurons AIA and ADE through the metabotropic receptor MGL-1 (***López-Cruz et al., 2019***). Notably, glutamatergic signaling alone was found not to be essential for this suppression, implying that additional factors contribute to the control of AIA/ADE output and, by extension, to the foraging transition (***López-Cruz et al., 2019***), a role that neuromodulators such as betaine are well placed to fill. Whether this transition reflects a discrete behavioral-state switch at all has recently been questioned: ***Margolis and Gordus (2025)*** show that a state-free stochastic model can reproduce the apparent local-to-global transition without invoking a switch, underscoring that the molecular and circuit basis of foraging dynamics remains actively debated. While LGC-41 is likely responsible for the inhibitory effects on local-search-promoting neurons, ACR-23 may mediate distinct, possibly excitatory functions depending on its expression context. By mapping the anatomical targets of betaine-associated neurons within the connectome, the Connectome Toolbox provides a starting point for future receptor-identification efforts. Combining these targets with expression data for LGC-41 and ACR-23 would separate candidates that could respond through known channels from those that would require as-yet unidentified receptors.

Taken together, these neurotransmitter overlays illustrate a central strength of the Connectome Toolbox — its ability to connect molecular identity (neurotransmitter and receptor expression), circuit architecture (the anatomical wiring they act upon), and behavior (foraging, locomotion, feeding) within a single framework.

### Using custom Python scripts to investigate bilateral symmetry in the *C. elegans* connectome

Bilateral symmetry is a fundamental organizational principle of the *C. elegans* nervous system and provides a biologically meaningful benchmark for comparative connectomics (***Brittin et al., 2021***; ***Birari and Rabinowitch, 2024***). Quantifying how this symmetry is preserved or altered across datasets, developmental stages, and connectivity modalities requires consistent identification of contralateral neuron pairs and uniform handling of heterogeneous connectome reconstructions, tasks that are typically non-trivial when working across independently published datasets.

To demonstrate how the Connectome Toolbox facilitates such analyses, we applied the bilateral symmetry index as defined by ***Kim et al. (2026)*** across multiple connectome datasets. This index quantifies bilateral symmetry in a connectome by measuring the proportion of connections that have a corresponding contralateral counterpart. Neurons are paired across the left–right axis, and each directed connection is classified as bilaterally symmetrical if an equivalent connection exists between the contralateral neuron pair (e.g. both ADLL→ASEL and ADLR→ASER connections are present). The resulting index ranges from 0 (fully asymmetric) to 1 (perfect bilateral symmetry), providing a quantitative measure of left–right structural correspondence while accommodating imperfect biological symmetry.

Our aim was not to re-evaluate the symmetry metric itself, but to illustrate how the toolbox enables reproducible, cross-dataset application of published analytical methods using a shared programmatic interface. A first step was to add an option to display the symmetry of individual connections for each dataset on the website. Visualizations of the individual connections that are bilaterally symmetrical and those that are not can be seen for two datasets at different developmental stages in Fig. 3, supplement 3a,b. While these views are useful for visualizing the presence or absence of individual connections and their bilateral counterparts, they do not provide a quantitative measure of overall symmetry across the entire connectome.

To illustrate the application of the Connectome Toolbox API, we computed symmetry indices across multiple chemical synaptic connectomes (***White et al., 1986***; ***Cook et al., 2019***; ***Witvliet et al., 2021***; ***Yim et al., 2024***) from different developmental stages (a screenshot of a Jupyter notebook containing the analysis is shown in Fig. 3, supplement 3c). Because all datasets are accessed through a common internal representation with standardized neuron identities and bilateral pairings, the same analysis could be applied without dataset-specific preprocessing. The overall level of bilateral symmetry across all neurons is broadly preserved across development (thick purple line, Fig. 3, supplement 3c). However, there are differences between the level of bilateral symmetry within subsets of somatic (non-pharyngeal) neurons, with sensory neurons generally less symmetrical and motorneurons generally more symmetrical, particularly in adult specimens. Most of these datasets focus on reconstructed neurons of the nerve ring and so include head and sublateral motorneurons but few or no ventral cord motorneurons (an exception to this being the relatively complete ***Cook et al. (2019)*** dataset). Ventral cord motorneurons (e.g. AS, DA, DB, DD, VA, VB, VD) are unpaired cells with somas along the the ventral midline. Under the symmetry index, each ventral cord motorneuron maps onto itself, so connections among them are classified as symmetrical by construction, raising the index independently of any biological left-right correspondence.

To help explore this, we used two of the ConnectomeViews in the Connectome Toolbox API, one containing all somatic motorneurons, and one containing only head/sublateral motorneurons. This allowed symmetry indices to be computed for these specific subsets, as well as for interneurons, sensory neurons and all neurons. For ***Cook et al. (2019)***, motorneuron symmetry is high when considering all somatic motorneurons (approx 0.9) but falls below 0.7 when only head/sublateral motorneurons are included (orange vs. dashed green line in Fig. 3, supplement 3c). In the other datasets, which contain few or no ventral cord motorneurons, the two views give near-identical values, as expected. The elevated motorneuron symmetry of ***Cook et al. (2019)*** therefore reflects differences in reconstruction coverage rather than biology.

This example demonstrates how the Connectome Toolbox enables published connectomic analyses to be easily run across datasets, modalities, and developmental contexts, with a small amount of scripting using the Connectome Toolbox API. By providing a unified and extensible framework for accessing and analyzing diverse connectome data, the toolbox lowers the barrier to comparative studies and supports reproducible exploration of structural principles such as bilateral symmetry.

### Integration of contact-based, neuropeptidergic, and functional connectivity

In addition to classical chemical synapses and electrical gap junctions, we have incorporated connectomic datasets containing physical connections (“contactome”), neuropeptidergic connections and functional connections. Contact-based connections, derived from nanoscale-resolution ultra-structural reconstructions, capture physical proximity between neurons independent of synaptic specialization (***Brittin et al., 2021***). A complete contactome provides a geometric and spatial framework of neuronal adjacency that can be used to examine organizational constraints underlying the distribution of chemical synapses and electrical gap junctions. The contactomes of ***Brittin et al. (2021)*** and ***Yim et al. (2024)*** (Fig. 2, supplement 1) have been integrated into the Connectome Toolbox, allowing users to explore how physical proximity relates to synaptic connectivity and to identify potential sites of non-synaptic interactions.

Neuropeptidergic connectivity represents extrasynaptic signaling relationships inferred from neuropeptide expression and receptor localization, capturing modulatory interactions that extend beyond anatomically defined synapses. Neuropeptide signaling can influence neuronal and circuit activity over broader spatial and temporal scales, providing a mechanism through which behavioral states and context-dependent modulation may be regulated in ways that are not readily explained by strictly anatomical synaptic connectivity alone (***Bargmann, 2012***; ***Marder and Thirumalai, 2002***; ***Ripoll-Sánchez et al., 2023***). Datasets of the preliminary neuropeptidergic connectome from ***Bentley et al. (2016)***, as well as the updated connectomes of ***Ripoll-Sánchez et al. (2023)*** which were generated using large-scale GPCR deorphanization, single-neuron scRNA-seq, and anatomical diffusion constraints (including short-, mid- and long-range diffusion models), have been incorporated into the framework (Fig. 4c). This allows users to compare these distinct versions, and explore how neuromodulatory interactions overlay onto the anatomical wiring diagram.

Functional connections, derived from direct perturbation of individual neurons, capture the sign, strength and directionality of one neuron’s causal influence on others (***Randi et al., 2023***). Unlike the correlational “functional connectivity” familiar from neuroimaging, these measurements are obtained by optogenetically activating a single identified neuron and recording the evoked responses across the rest of the network, yielding a directed, causal map of signal propagation. Such perturbation-based measurements reveal network-level coordination and extrasynaptic signaling that are not fully accounted for by static anatomical connectivity alone (***Randi et al., 2023***). The functional connectivity data from ***Randi et al. (2023)*** we have incorporated can be visualized using the same tools as anatomical connectivity, e.g. plotting the relative excitatory or inhibitory effect on all other neurons when a single identified neuron is optogenetically stimulated (Fig. 4d).

### Comparing computational models of worm locomotion to connectivity datasets

Computational models of *C. elegans* neuromechanics have become important tools for investigating how neural circuits generate behavior, particularly locomotion (***Cohen and Denham, 2019***). By making the assumptions of a theory explicit and testing whether they are sufficient to reproduce observed behavior, such models offer a way to compare competing mechanistic hypotheses, e.g. whether the rhythmic bending waves of locomotion are driven principally by intrinsic network central pattern generators along the ventral nerve cord (***Olivares et al., 2018***; ***Izquierdo and Beer, 2018***) or by proprioceptive stretch-receptor feedback coupling successive body segments (***Boyle et al., 2012***; ***Wen et al., 2012***), or by some combination of the two.

Although these models aim to reproduce the dynamics of locomotion, they necessarily simplify underlying anatomical connectivity, complicating direct comparison with empirical anatomical datasets. As an example application of the Connectome Toolbox, we compared the network components of published neuromechanical models of *C. elegans* with the anatomical connectome of ***Cook et al. (2019)***, coupled with the neurotransmitter identity information from ***Wang et al. (2024)***. We used two models that simulate the dynamical activity of neural circuitry and muscle dynamics during locomotion (***Gleeson et al., 2018***; ***Olivares et al., 2021***), and using the toolbox, we generated matched graphical representations of locomotion-related circuitry from each model (Fig. 8d–i) and from the anatomical dataset (Fig. 8a–c), enabling direct comparison of chemical connections and electrical connectivity in the context of empirical anatomy. Cholinergic and GABAergic connections are selected in the anatomical dataset (for putative excitatory and inhibitory connections, respectively), which can be compared to the patterns of positive and negative connections in the models. Connectivity matrices showing the structured connectivity within and between motorneurons and muscles are shown in Fig. 8, supplement 1.

**Figure 8.**
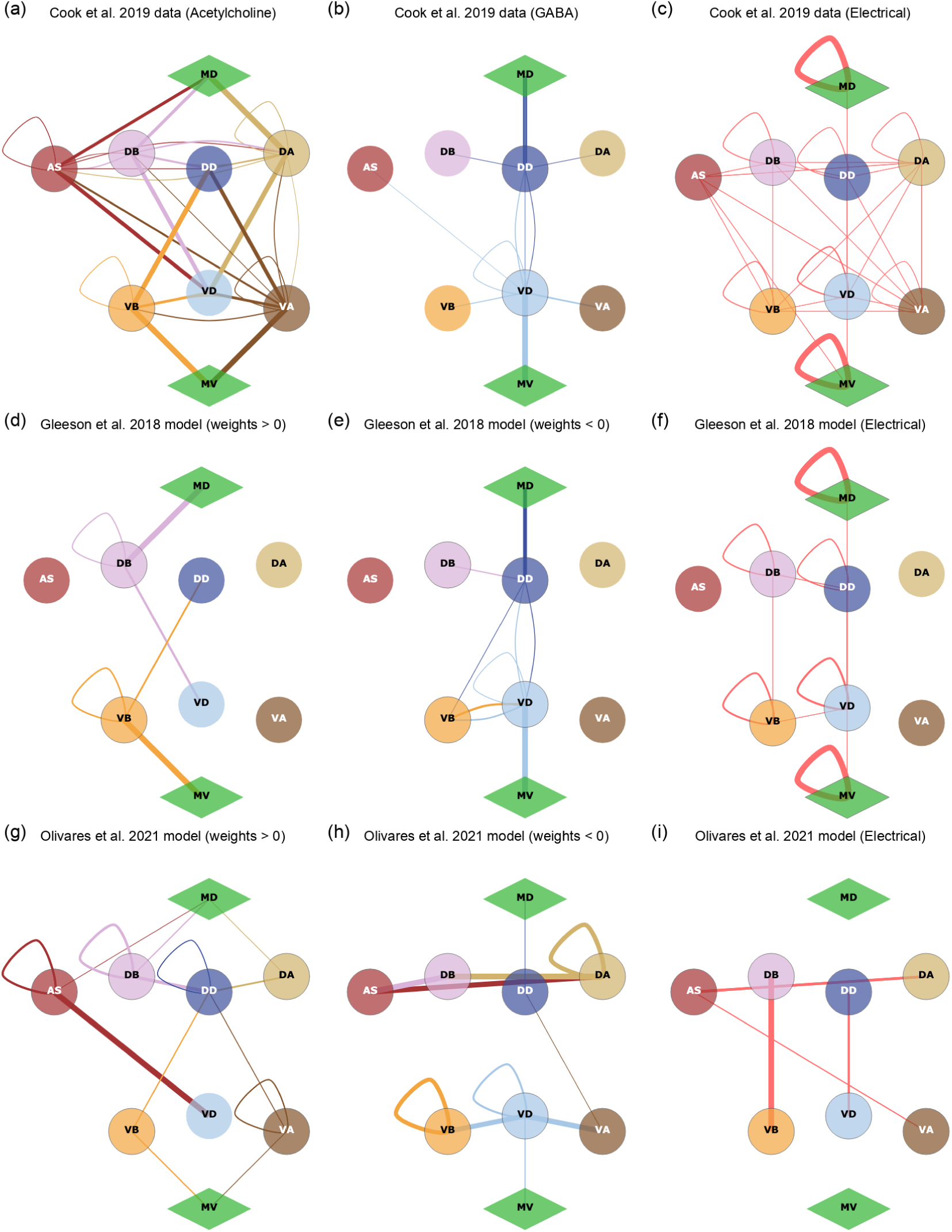
Comparing network connectivity in computational models of *C. elegans* locomotion (***Gleeson et al., 2018***; ***Olivares et al., 2021***) with experimental connectome data (***Cook et al., 2019***; ***Wang et al., 2024***). A connectome view consisting of the main ventral cord motorneurons and muscles involved in forward and reverse locomotion is shown for three different networks. The nodes consist of dorsal (D*) and ventral (V*) body wall muscles (MD, MV) as well as B-type (DB, VB), A-type (DA, VA) and D-type (DD, VD) motorneurons, and AS-type motorneurons which mainly target dorsal muscles. (a-c) Anatomical connectivity from ***Cook et al. (2019)***, coupled with neurotransmitter identities from ***Wang et al. (2024)***. (a) All chemical connections from cells releasing acetylcholine (putatively excitatory) according to ***Wang et al. (2024)***. (b) All chemical connections from cells releasing GABA (putatively inhibitory) according to ***Wang et al. (2024)***. (c) All electrical connections. (d-f) The connectivity used in the ***Gleeson et al. (2018)*** computational model. (d) All chemical connections with a positive weight, (e) with a negative weight, and (f) all electrical connections. (g-i) The connectivity used in our selected instance from the ***Olivares et al. (2021)*** computational model. (g) All chemical connections with a positive weight, (h) with a negative weight, and (i) all electrical connections. **Figure 8–F**igure supplement 1. Adjacency matrix based views of these connections.

Importing the network structure from the computational models into Connectome Toolbox was facilitated by the use of the NeuroML standard (***Cannon et al., 2014***; ***Sinha et al., 2025***), which allows specification of structured representations of neuronal and synaptic properties, as well as network connectivity. We developed a DataReader class specifically to parse model files in this format containing *C. elegans* circuitry, extracting the relevant connectivity information and mapping it onto the standardized neuron identities used in the Connectome Toolbox. The model of ***Gleeson et al. (2018)*** is natively built in NeuroML (using the c302 framework), while the C++ based model code from ***Olivares et al. (2021)*** was extended to export the relevant connectivity data in NeuroML format (see Methods).

### Gleeson et al. 2018 model

***Gleeson et al. (2018)*** described the c302 framework, which can be used to generate computational models of *C. elegans* neuromuscular circuitry at multiple scales and levels of biological detail. To demonstrate the utility of the package, a simple model was generated which reproduced key qualitative features of the real *C. elegans* locomotion circuit, such as head-to-tail traveling waves of muscle activation consistent with experimental observations. However, this incorporated a number of simplifying assumptions, meaning the model reflects the worm’s circuitry at a coarse, functional level rather than as a fully data-constrained, biologically complete connectome implementation.

The model incorporates bilateral AVB command neurons, 18 B-type motorneurons, and 19 inhibitory D-type motorneurons, with wiring data which were obtained from the WormWiring project at the time (which consisted of a near-complete version of the ***Cook et al. (2019)*** connectome). All synaptic connections were assigned equal weight, despite variation in the number of individual synaptic contacts in the anatomical connectome. To approximate central pattern generator activity during forward locomotion, rhythmic current pulses were applied to B-type motorneurons and muscles near the head. In addition, proprioceptive feedback to posterior motorneurons was represented by an effective excitatory mechanism directed from each B-type motorneuron to its posterior neighbor.

When viewed through the Connectome Toolbox, overlaying neurotransmitter identity and connection class information shows that the majority of cholinergic (putative excitatory) chemical connections between the populations included in the ***Gleeson et al. (2018)*** model are present in the anatomical connectome (Fig. 8a,d). Note, populations which are not included in the model (AS, DA, VA), and therefore have no connections, are still shown here, see below. The overall structure of GABAergic/inhibitory connectivity is preserved (Fig. 8b,e). However, this model explicitly included inhibitory connections from B-type to D-type motorneurons on both the dorsal and ventral sides, to facilitate oscillatory activity leading to forward locomotion. These are not reflected in the GABAergic connectivity of the ***Cook et al. (2019)*** dataset (Fig. 8b). While inhibition can take place through glutamate-gated chloride channels (***Horoszok et al., 2001***; ***Sato et al., 2021***), investigation of the ***Wang et al. (2024)*** neurotransmitter data indicates that these neurons are primarily cholinergic.

Fig. 8, supplement 1c and f show that the overall pattern of observed electrical connectivity, chiefly connections between adjacent B-type and D-type motorneurons as well as between adjacent muscles in each quadrant, are incorporated in the model. The graphical comparison between the anatomical dataset and the ***Gleeson et al. (2018)*** model (Fig. 8a-f) also makes it explicit that AS, DA, and VA motorneuron classes are not included in the modeled circuitry, as no connections to/from them are present in the model. DA and VA motorneurons are involved in reverse locomotion (***Haspel et al., 2010***), which the model was not designed to simulate, and AS neurons, which are involved in forward locomotion (***Tolstenkov et al., 2018***), were not required here.

### Olivares et al. 2021 model

Olivares and colleagues developed a neuromechanical model that integrates neural connectivity with muscle activity and body mechanics (***Olivares et al., 2021***). The model consists of seven identical repeating subunits, each representing a segment of the ventral nerve cord, connected to a previously published 2D model of the worm body (***Boyle et al., 2012***). Each subunit contains seven classes of motorneurons (AS, DA, DB, DD, VA, VB, VD), with individual motorneurons mapped to corresponding muscle segments based on a statistical analysis of motorneuron–muscle positioning (***Haspel and O’Donovan, 2011***). This model extends earlier work by the same group (***Olivares et al., 2018***; ***Izquierdo and Beer, 2018***). The neuromechanical model was designed to investigate the hypothesis that a chain of multiple network rhythmic pattern generators in the ventral nerve cord can drive forward locomotion, without requiring stretch-receptor feedback. Multiple configurations of the model were generated by an evolutionary algorithm, and an analysis was then performed to identify those that produced realistic locomotion kinematics.

Fig. 8g-i and Fig. 8, supplement 1g-i show graphical representations of a single instance of the model, which was selected for its ability to produce realistic forward locomotion (***Olivares et al. (2021)*** Fig. 4). Fig. 8g-i show that the model explicitly incorporates AS, DA, and VA motorneuron classes, resulting in a circuit structure that differs from the ***Gleeson et al. (2018)*** model and more closely reflects the diversity of motorneuron types present in the anatomical connectome. AS neurons play a prominent role in driving forward locomotion (***Tolstenkov et al., 2018***), and significant cholinergic connections from these are present in the anatomical dataset and form a core part of the excitatory (positive weight) motor circuitry of the model (Fig. 8g). The model includes inhibitory/negative weight connections such as DB→AS and AS→DA (Fig. 8h), which are not present in the reconstructed GABAergic dataset. This is a consequence of the sign and strength of connections between motorneurons being unconstrained during the evolution of the model, to better examine the space of possible circuit configurations that can produce realistic locomotion. Gap junction connections are present in the model also, but the overall pattern of electrical connectivity is less similar to the anatomical dataset than in the ***Gleeson et al. (2018)*** model, and the illustrated network doesn’t include connections between adjacent B-type and D-type motorneurons or between adjacent muscles, which the modelers explicitly excluded from their model (Fig. 8, supplement 1f,i). While this is just a single instance of the networks produced by the evolutionary algorithm in ***Olivares et al. (2021)***, any of the other generated configurations could be similarly visualized, and quickly compared to the known experimental data.

These comparisons illustrate how the Connectome Toolbox enables systematic evaluation of computational model architectures against anatomical connectivity, facilitating investigation of the biological plausibility and structural assumptions underlying neuromechanical models of locomotion.

## Discussion

The journey toward a comprehensive connectome of *C. elegans* remains incomplete. Early optimism that a static wiring diagram alone could explain the worm’s neural functions has proven overly simplistic. The synaptic connectome captured from electron microscopy is a snapshot of an organism’s nervous system that, in reality, exhibits plasticity, neuromodulation, and individual variability. Key modes of neuronal communication, such as extrasynaptic monoamine and neuropeptide signaling, fall outside the scope of a traditional synapse-only map. In this work, we address these gaps by consolidating multiple modes of connectivity data, produced by different labs and covering many developmental stages into a single framework. This integrative Connectome Toolbox provides a clearer framing of how various datasets interrelate, and represents a novel open-science infrastructure to accelerate connectomics research. By unifying anatomical, extrasynaptic, and functional connectivity information, our platform facilitates reproducible analyses and hypothesis generation that were previously hindered by fragmented data. It is a community resource explicitly designed for extensibility, so it can evolve as new datasets emerge, ultimately providing a comprehensive set of multimodal connectome datasets, compliant to the FAIR data sharing principles (***Wilkinson et al., 2016***) and ready for integration into multiscale models of the worm nervous system, that can be used to investigate *C. elegans* behavior *in silico* with high fidelity.

Among the example analyses presented, the integration of betaine-associated neurons into the anatomical connectome (Fig. 7h) highlights a particularly promising application of the multimodal framework. Betaine has recently been identified as an inhibitory neuromodulator critical for mediating the switch from local to global foraging behavior (***Hardege et al., 2022***), yet the full set of betaine-responsive cells remains to be characterized. By overlaying the expression of betaine-handling machinery (snf-3/BGT1 uptake and cat-1/VMAT vesicular loading) from ***Wang et al. (2024)*** onto the connectome, we mapped the anatomical postsynaptic partners of neurons with the capacity to take up and potentially release betaine. Because this overlay does not encode receptor expression, it does not show that these partners respond to betaine. Instead, it defines a candidate set of cells that could be cross-referenced with expression data for the known betaine-gated channels LGC-41 and ACR-23. Candidates that lack both channels yet receive input from betaine-associated neurons would be priority targets in the search for additional betaine receptors. This illustrates how the Connectome Toolbox can move beyond data consolidation to generate testable hypotheses about circuit function.

### Validation, extensibility and usage of the Connectome Toolbox

To demonstrate the validity, flexibility, and practical utility of the Connectome Toolbox, we applied it across multiple datasets and use cases, including direct comparison of experimentally reconstructed connectomes with computational models of *C. elegans* locomotion. As shown in Fig. 8, the toolbox enables consistent representation and comparison of anatomical, molecular, and model-derived connectivity, making explicit how empirical wiring, neurotransmitter identity, and modeling abstractions relate within a unified framework.

While there has been much interest in using the *C. elegans* connectome outside of the field for graph theoretical analysis etc., it is not often clear which exact version of the connectome has been used. ***White et al. (1986)*** and ***Cook et al. (2019)*** are the most widely cited publications, but even in these cases, the former did not release a structured dataset with that publication, and the latter’s data has been updated with corrections since publication (https://wormwiring.org/pages/ adjacency.html). ***Supplementary file 1*** documents the curation process we have gone through in retrieving, testing and standardizing each of the datasets. It also describes the suite of validation tests included with the toolbox which ensures that the data retrieved through our API accurately reflect the contents of the source files of the datasets.

### Usage across levels of technical expertise

The Connectome Toolbox supports multiple modes of interaction, reflecting the diversity of the *C. elegans* research community (Fig. 2). Researchers without programming experience can explore connectome datasets through the public web interface (https://openworm.org/ConnectomeToolbox), which provides pre-generated, interactive visualizations of connectivity across modalities, developmental stages, and sexes, as well as neuron-specific summaries (Fig. 2, supplement 1,2). Indeed, all of the panels in Fig. 4, Fig. 5, Fig. 6, Fig. 7, and Fig. 8 are available to explore interactively on the website, without requiring any programming knowledge.

Users with basic Python experience can install the Python package and use the provided Jupyter notebooks to programmatically access datasets, generate customized visualizations, and perform comparative analyses in a reproducible manner (Fig. 2, supplement 3; Fig. 3, supplement 3c). Advanced users can access and contribute to the open-source codebase, or use the package as a library, on which to build other software applications or online resources for *C. elegans* research.

### Extensibility and community contribution

The toolbox is designed as an extensible framework rather than a fixed dataset. The codebase is fully open source and so it encourages community contribution, review, and extension as new datasets and analysis methods become available.

Three specific extension types are envisioned which can easily be incorporated into the code-base by interested users: 1) new connectome datasets, 2) new visualization methods, and 3) new circuit level views. New connectome datasets can be incorporated by implementing a DataReader class that maps source-specific formats onto the common internal ConnectomeDataset representation. All existing visualization methods are designed to operate on this internal representation, so new datasets that conform to the standard format can be immediately visualized and compared with existing datasets. Current visualization methods are based on the plotly package for interactive graphing, and new interactive graphs, etc. can be added by implementing a method in the ConnectomeDataset to return a plotly figure. Finally, new circuit level views can be implemented by creating a ConnectomeView class with nodes positioned in 2D, each containing sets of cells to represent a specific anatomical circuit (see Methods).

These design choices ensure that the toolbox can evolve with the field, incorporating new data and methods, and encouraging contributions from users with different interests/skill sets, without requiring fundamental changes to the underlying architecture.

### Relation to OpenWorm project

The *C. elegans* Connectome Toolbox is an integral component of the larger OpenWorm open-science initiative (***Sarma et al., 2018***; ***Szigeti et al., 2014***), and has been designed as a core part of a unified ecosystem aiming to digitally reconstruct *C. elegans*. Specifically, it underpins the neural data layer of the OpenWorm project, and will supply curated connectome data to other tools and models.

The c302 simulation framework for generating computational models of *C. elegans* (***Gleeson et al., 2018***) has already incorporated the Connectome Toolbox as a library, to provide updated neuron lists and connectivity. This tight integration means that extensions in the toolbox, such as the addition of new connectome datasets, are easily available to simulations of worm behavior running in OpenWorm’s pipeline. Conversely, feedback from modeling efforts (e.g., identifying requirements such as the need for information on where synapses are located on dendrites) can guide toolbox development, creating a virtuous cycle of refinement.

Moreover, the toolbox embodies OpenWorm’s philosophy of openness and reproducibility (***Wilkinson et al., 2016***; ***Szigeti et al., 2014***). All data within it can be traced back to published sources or community contributions, and by centralizing these data with proper citations, validation tests, and version control, the toolbox helps ensure transparency and trust in derived works. Rather than each laboratory curating its own version of the connectome, the community can converge on this shared resource, reducing duplication of effort. This resource is designed to complement other OpenWorm components: both the Sibernetic physics engine used for 3D worm body simulations (***Palyanov et al., 2018***), as well as extensions to 2D neuromechanical models (***Olivares et al., 2021***; ***Izquierdo and Beer, 2018***) will require testing with the most up-to-date connectivity data for neurons and muscles, and the toolbox provides a standardized interface for this purpose.

Together, these components advance the project toward a comprehensive *in silico* worm by making multimodal *C. elegans* connectome data ready for integration into multiscale whole-organism models.

### Relationship to other ***C. elegans*** neuroinformatics resources

Beyond its role within OpenWorm, the Connectome Toolbox exists within a broader ecosystem of *C. elegans* neuroinformatics resources, and is designed to build upon and operate alongside them rather than to supersede them. Curated knowledge bases such as WormAtlas (***Altun et al., 2025***) and WormBase (***Harris et al., 2020***) remain the authoritative references for cellular anatomy, nomenclature, and gene-level annotation, and we adopt their conventions directly, for example, the WormAtlas color scheme used throughout our visualizations. Likewise, WormWiring (***Cook et al., 2019***) and NemaNode (***Witvliet et al., 2021***) provide invaluable, purpose-built interfaces to the anatomical reconstructions they host.

More recent tools such as WormNeuroAtlas (***Randi et al., 2023***) offer programmatic access to specific datasets. The Connectome Toolbox builds on this directly, using the WormNeuroAtlas Python API to incorporate the functional connectivity data of ***Randi et al. (2023)*** (***Supplementary file 1***).

Complementary efforts are also emerging to integrate multimodal *C. elegans* data within unified computational environments. Notably, the recently described CeDNe (***Moza and Zhang, 2025***) organizes anatomical, molecular, and functional data into a multi-relational graph and couples this representation to built-in top-down dynamical modeling, using it to identify higher-order wiring motifs such as chained feed-forward loops. CeDNe represents a valuable addition to the field, and the two resources are complementary in emphasis: CeDNe focuses on graph-based analysis and the construction of dynamical models within a single environment, whereas the Connectome Toolbox prioritizes the reconciliation of many independently published datasets, spanning anatomical, contact-based, extrasynaptic, neurotransmitter, and functional modalities, onto a common annotation reference, with automated data visualization and analysis for non-computational researchers, and direct comparison of published computational models (via NeuroML) against anatomical data. Together, such frameworks reflect a broader movement toward integrative, reproducible connectomic infrastructure for *C. elegans* and, increasingly, for larger nervous systems.

### Limitations of current work

While we have made efforts to consolidate and standardize connectome datasets, there are inherent limitations in the underlying data that must be acknowledged. The original electron microscopy reconstructions, while groundbreaking, are based on a limited number of animals and may not capture the full extent of biological variability in the *C. elegans* nervous system (***White et al., 1986***). Subsequent reconstructions have attempted to address this by including additional specimens and developmental stages (***Witvliet et al., 2021***; ***Yim et al., 2024***), but the extent of variability across individuals, sexes, and conditions remains incompletely characterized. The datasets we have integrated are influenced by variations in reconstruction methodologies, annotation/metadata standards, and biological scope, which can introduce discrepancies that do not necessarily reflect true biological differences. While we have standardized neuron identifiers and naming conventions to mitigate this, some ambiguity may still exist, particularly for neurons with complex morphologies or those that are difficult to classify. Comparisons between datasets obtained by the same lab or consortium, using consistent methods (e.g. ***Witvliet et al. (2021)***), will always be more insightful than comparisons between datasets obtained by different labs using different methods. The sensory/inter/motorneuron trichotomy we adopt from ***Cook et al. (2019)*** is itself a simplification, as many *C. elegans* neurons are multifunctional; over half have demonstrated or possible sensory function, and several cells classified as motorneurons act as integrative hubs (***Emmons, 2024***), and the groupings used here should be read as a practical convention rather than a strict functional partition.

There are many relevant datasets which have not been included in the current release of the toolbox, but are candidates to be added in the future. For example, neuron-specific innexin expression data (***Altun et al., 2009***; ***Bhattacharya et al., 2019***) would help refine the electrical connectome.

Another crucial shortcoming in our current framework is the lack of data and visualizations related to the spatial distribution of synapses on neuronal processes. Such spatial information is critical for understanding how synaptic inputs are integrated by neurons, and how circuit motifs are organized in three-dimensional space. ***Ruach et al. (2023)*** illustrate the value of this: using the synaptic coordinates accompanying the ***Cook et al. (2019)*** reconstruction together with the ***Brittin et al. (2021)*** contactome, they showed that chemical synapses cluster along neurites far more than expected from anatomy alone, suggesting compartmentalized computations that an adjacency-matrix representation cannot capture. Volumetric reconstructions, as well as skeletonized 3D representations, are now available for the *C. elegans* nervous system (***Witvliet et al., 2021***; ***Brittin et al., 2021***; ***Yim et al., 2024***), and tools are beginning to make these spatial datasets more accessible; for example, NeuroSC (***Koonce et al., 2025***) provides an interactive 3D viewer for exploring neuron morphologies, membrane contacts, and synaptic positions across the developmental EM series.

We plan to extend the Connectome Toolbox with further online visualization tools which can be used to explore these types of datasets, alongside the other 3D data being used in the Open-Worm project (e.g. model neurons (***Gleeson et al., 2018***), worm body models (***Palyanov et al., 2018***), neuron specific fluorescent marker expression data from ***Yemini et al. (2021)***).

### Conclusion and vision for the future

The consolidation of multimodal connectivity data for *C. elegans* presented here provides a foundation for a more integrated view of how structure and signaling converge to produce behavior in a complete nervous system. The neurotransmitter overlays presented above illustrate this directly, connecting molecular identity, circuit wiring, and behavior within one framework. By unifying anatomical, extrasynaptic, and functional datasets into a single extensible framework, the *C. elegans* Connectome Toolbox offers a model for how connectomic resources can evolve from static wiring diagrams into dynamic, community-driven platforms for data integration and discovery. Additionally, because every value in our toolbox is curated, versioned and traceable to a published source, it is also well suited to serve as a grounding layer for AI applications such as Large Language Model (LLM) based assistants for *C. elegans* research, where answers drawn from a validated, provenance-tracked dataset can be more trustworthy and verifiable than those generated from unstructured text alone.

While *C. elegans* remains the only organism whose complete nervous system has been reconstructed at cellular resolution across sexes and developmental stages, the conceptual and computational challenges of synthesizing diverse data modalities are now being confronted in higher-order systems. Recent large-scale connectomics projects in Drosophila melanogaster (***Scheffer et al., 2020***; ***Lin et al., 2024***) and in the mouse brain (***Yao et al., 2023***; ***MICrONS Consortium, 2025***) exemplify the accelerating scale and complexity of efforts to reconstruct complete brains at synaptic or cell-type resolution. The framework we present here addresses precisely the types of integration and standardization problems that will become increasingly critical for these larger systems. The need to harmonize anatomical, molecular, and functional information across modalities and scales is not unique to *C. elegans*, it is a defining challenge for all of neuroscience.

In this sense, the work described here provides a logical and methodological bridge between the historical completeness of the *C. elegans* connectome and the emerging landscape of vertebrate and insect brain mapping. The same principles (open data access, consistent data formats, and flexible multimodal analysis pipelines) will be essential for the synthesis of next-generation connectomes. As datasets for Drosophila and rodent brains become richer and more multimodal, the lessons learned from building and maintaining a fully integrated connectomic framework at the scale of *C. elegans* will remain indispensable for realizing a unified, multiscale understanding of brain architecture across species.

Looking forward, frameworks such as the *C. elegans* Connectome Toolbox can help transform connectomics from an archival science into a predictive one. By linking structural, molecular, and functional datasets within a computationally tractable organism, this work paves the way for integrative models capable of simulating emergent behavior directly from data. Such simulation-ready frameworks will ultimately serve as a proving ground for the methodologies needed to interpret larger, more complex connectomes, from Drosophila to rodents, species which are also the subject of whole body simulation efforts (***Lobato-Rios et al., 2022***; ***Aldarondo et al., 2024***; ***Vaxenburg et al., 2025***). In this way, the worm continues to illuminate the path toward a mechanistic understanding of nervous systems.

## Materials and Methods

### Software implementation and availability

The *C. elegans* Connectome Toolbox is implemented as an open source Python package, cect, which is being developed at (https://github.com/openworm/ConnectomeToolbox) and is compatible with Python version 3.9+. A high level overview of the architecture is shown in Fig. 2. Core dependencies include NetworkX (https://networkx.org) for graph representation and analysis and plotly (https://plotly.com) for interactive visualization; a full dependency list is given in the repository. The package can be installed from PyPI (using: pip install cect).

Example Jupyter notebooks demonstrating the use of the Python API for data exploration, analysis, and visualization (see Fig. 2, supplement 3 and Fig. 3, supplement 3c) are included in the repository (docs/notebooks directory) and available to browse online, e.g. https://openworm.org/ConnectomeToolbox/notebooks/Introduction.

### Data sources, provenance and validation

Connectivity datasets were obtained from the original published sources, or community resources such as WormWiring, in their native formats (text, Excel/OpenOffice spreadsheets, CSV, JSON). All incorporated datasets, along with the original publications from which they came, short descriptions, and the interpretation of weights between connections in each dataset, are listed in Table 1. These entries also include links to a webpage with a representative view of an adjacency matrix for the dataset, as well as to the online validation test pages, which include full details of where the original datasets were obtained. ***Supplementary file 1*** describes the curation process for each dataset, as well as the current validation test results.

### Standardization: DataReader and ConnectomeDataset classes

Each source dataset is parsed by a dedicated DataReader class that converts the original file format into a common internal representation, the ConnectomeDataset, which stores the list of cells present and independent adjacency matrices for each connection class (e.g. chemical, electrical). Cell names for neurons, muscles, and support cells are standardized onto the reference nomen-clature of ***Cook et al. (2019)***; neurons are classified as sensory, interneurons, or motorneurons following the same reference.

### ConnectomeView definitions for analysis and visualization

ConnectomeView classes define subsets of cells or grouped nodes used for visualization and analysis (e.g. pharyngeal neurons; sensory/inter/motorneuron groupings; the escape-response circuit; the ***Cook et al. (2019)*** Fig. 3–style whole-network layout). When converting a ConnectomeDataset to a ConnectomeView, all the existing connections between cells in node A and cells in node B are summed, to produce a single weight of the connection from node A to node B.

A textual description can be added to each node, and additionally a preferred color, an (x,y) position, a shape (e.g. circle, triangle), and preferred size can be set which can be used across graphical (and textual) depictions for a consistent representation of the view (see Fig. 7 and Fig. 7, supplement 1).

### Visualization methods

All views can be rendered as one or more graphical types. Adjacency matrices (Fig. 4) are drawn as 2D heatmaps with cells grouped by type and colored by WormAtlas convention (Appendix 1 Fig. 1); cell values depict connection weights (as described in Table 1).

Graph based views either render the nodes in the current view as fixed locations in 2D, if there are (x,y) position associated with the nodes of the ConnectomeView (as in Fig. 7, Fig. 8), or use force-directed layouts generated by the NetworkX package for laying out the nodes (Fig. 5).

Hive plots (Fig. 6) assign nodes to fixed axes by neuron class (six axes: two per sensory/interneuron/motor type), position nodes by total degree (incoming + outgoing), and color edges by presynaptic neuron type, with internal connections in gray. Python package hiveplotlib is used to generate these visualizations.

All of these graphical views can be rendered as static images (PNG) or interactive plotly figures (HTML) for use online or in Jupyter notebooks.

### Website generation

The public web interface (https://openworm.org/ConnectomeToolbox) presents pre-generated, interactive views produced from the same internal ConnectomeDataset/ConnectomeView structures. The website is built using the static site generator MkDocs (https://www.mkdocs.org) (Material theme), with custom JavaScript to embed the interactive plotly figures. It is hosted at GitHub Pages, with commits to the main branch automatically triggering a rebuild and redeployment of the site, dependent on a number of automated checks passing, including all validation tests on datasets.

### Import of computational models using NeuroML

Computational models were imported via a dedicated DataReader that parses circuitry described in a file in the NeuroML standard (***Cannon et al., 2014***; ***Sinha et al., 2025***). Populations in the NeuroML model can either contain a single cell, with a population id matching a known cell type (e.g. id=“ADAL”, id=“AS2”, each with one cell), or can contain a list of cells with the population id appended to the position in the list (not index, which starts at 0) to create a known cell id (e.g. id=“AS”, containing 10 cells with mapping AS[0] -> AS1, AS[1] -> AS2, etc.). Projections in the NeuroML model are mapped to connections in the ConnectomeDataset, according to synapse type.

The c302 model from ***Gleeson et al. (2018)*** was natively expressed in NeuroML, and the file https://github.com/openworm/c302/blob/master/examples/c302_C2_FW.net.nml (i.e. C2 parameter set (conductance based point neurons, analogue synapses), FW configuration for forward locomotion) was used. Note the C1 configuration was used in the original paper, but this was a simplified version of the C2 configuration. Both configurations have the same network structure (hence identical connectivity diagrams), but the latter has been used more extensively in the OpenWorm project, in particular for driving activity of the 3D worm body simulation using the Sibernetic simulator (***Palyanov et al. (2018)***, https://github.com/openworm/OpenWorm).

The GitHub repository containing the C++ code for the ***Olivares et al. (2021)*** model contained all of the optimized models (phenotypes) generated by the evolutionary algorithm outlined in that paper. We created a fork of this repository, and added a Python script to parse the generated optimized model instances and convert them into NeuroML format: https://github.com/openworm/MultipleNetworkOscillator/blob/master/nml_musc_generator.py. The specific instance of the model used in Fig. 4 in that paper had been labeled 23, and the generated NeuroML file for this instance is available at https://github.com/openworm/MultipleNetworkOscillator/blob/master/Filtered_Solutions_fig_2_3_4_5_6/filtered_agents/23/Worm2D_musc.generated.net.nml.

## Acknowledgments

We would like to thank the many members of the OpenWorm community who have contributed to, and helped test, the Connectome Toolbox package, as well as the various software packages which have led up to its development, in particular Mark Watts for the owmeta/PyOpenWorm project. We thank the OpenWorm Foundation for providing community infrastructure supporting the development and dissemination of this resource. We also thank the broader *C. elegans* research community for their feedback and suggestions on the design and functionality of the Toolbox, and for their efforts in generating and sharing the datasets that form its core content.

## Additional information

### Code availability

Source code is available at https://github.com/openworm/ConnectomeToolbox under the MIT License. The version used for the analyses in this manuscript is release v0.3.5, archived at https://github.com/openworm/ConnectomeToolbox/releases/tag/v0.3.5.

### Data availability

All datasets integrated into the Toolbox can be traced back to previously published sources; no new primary data were generated in this study. The datasets are located in the cect/data directory of the repository, most of which are identical to the supplementary information released with the original publications (https://github.com/openworm/ConnectomeToolbox/tree/main/cect/data). ***Supplementary file 1*** gives full information on how these datasets were curated, and the very latest validation information is available at https://openworm.org/ConnectomeToolbox/Validation.

## Funding

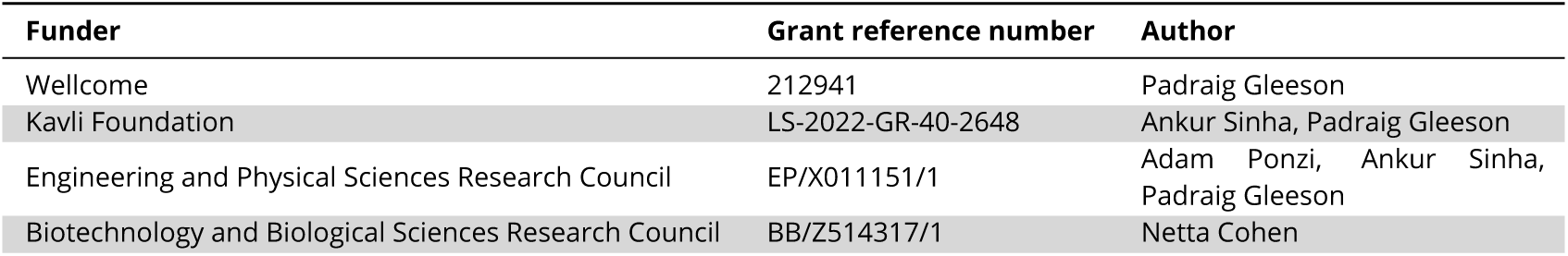

## Additional files

### Supplementary files

***Supplementary file 1***. Describes the suite of tests used to validate the data readers in the Connectome Toolbox, and provides information on the original data sources, any issues found with these data, and how they were addressed.

**Appendix 1 Figure 1.**
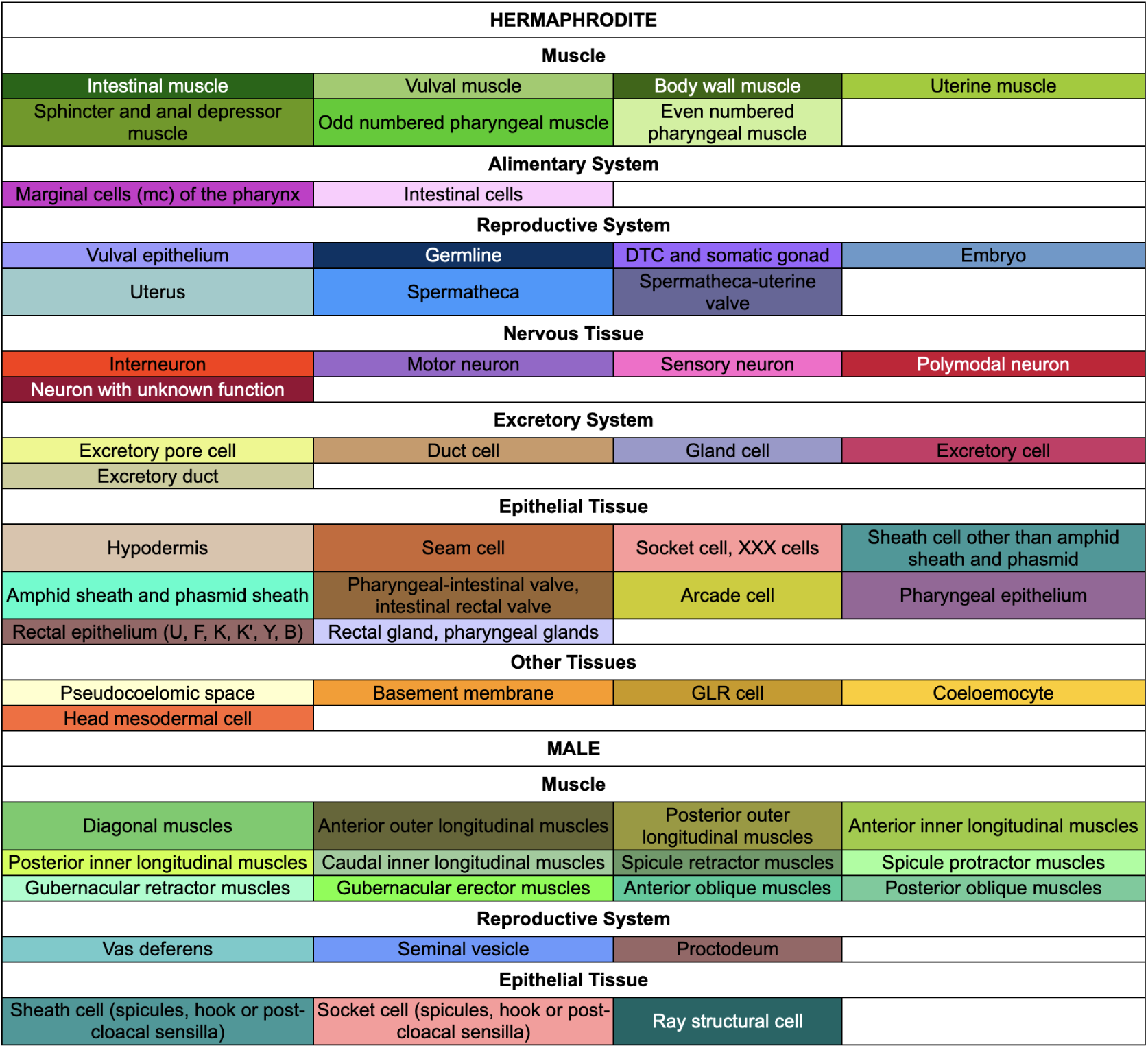
Colors used for cell types in Fig. 3 and Fig. 5, as well as axis labels in Fig. 4 and cells/labels in Fig. 6. Colors are based on those used across WormAtlas, see https://www.wormatlas.org/colorcode.htm.

---

**Figure 2–Figure supplement 1.**
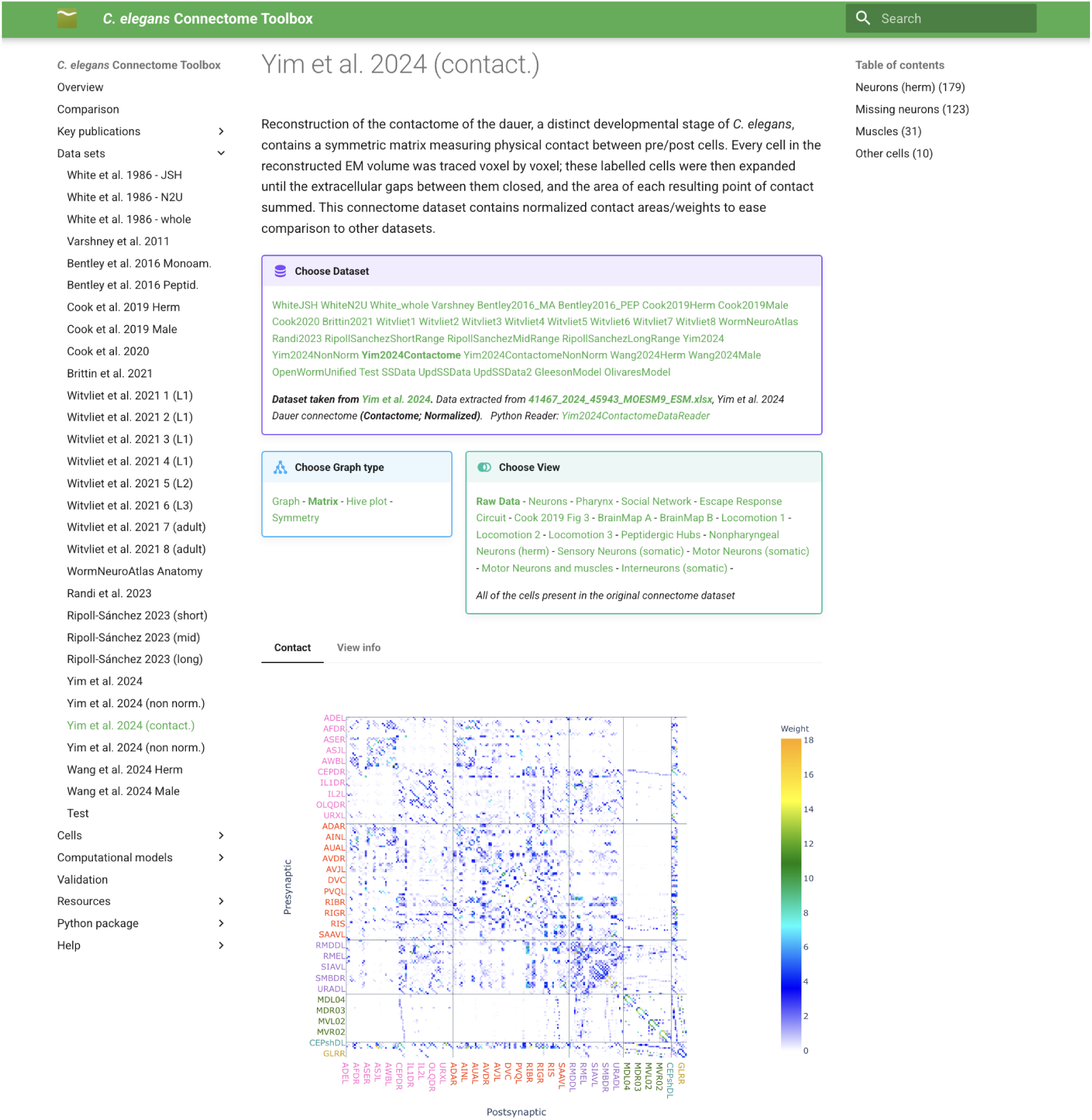
Screenshot of a page on the Connectome Toolbox website for a single dataset—the dauer contactome from ***Yim et al. (2024)***. Hyperlinks in the three boxes at the top allow quick switching between the datasets, changing the graph type from matrix to hive plot, etc., as well as selecting the view for subsets of cells or circuits constructed from the connectome data. An adjacency matrix view of the measured physical contacts between cells in the dataset is shown on the bottom. As expected for contactome data, the matrix is symmetric.

**Figure 2–Figure supplement 2.**
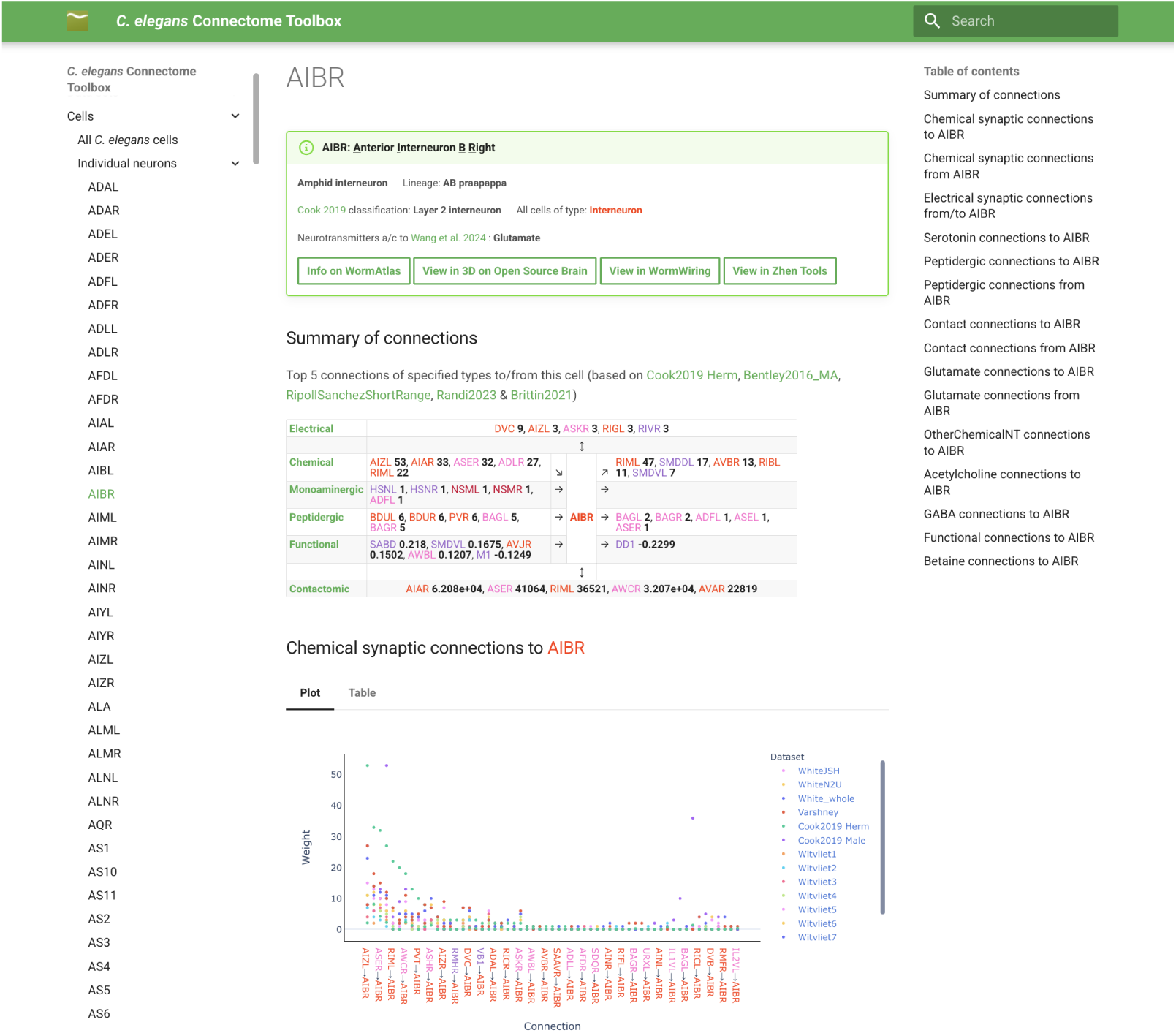
Screenshot of a page on the Connectome Toolbox website for a single cell, AIBR. Provides information on the cell type, neurotransmitter identity (if known), and connectivity information (of multiple modalities) to/from other cells in the dataset.

**Figure 2–Figure supplement 3.**
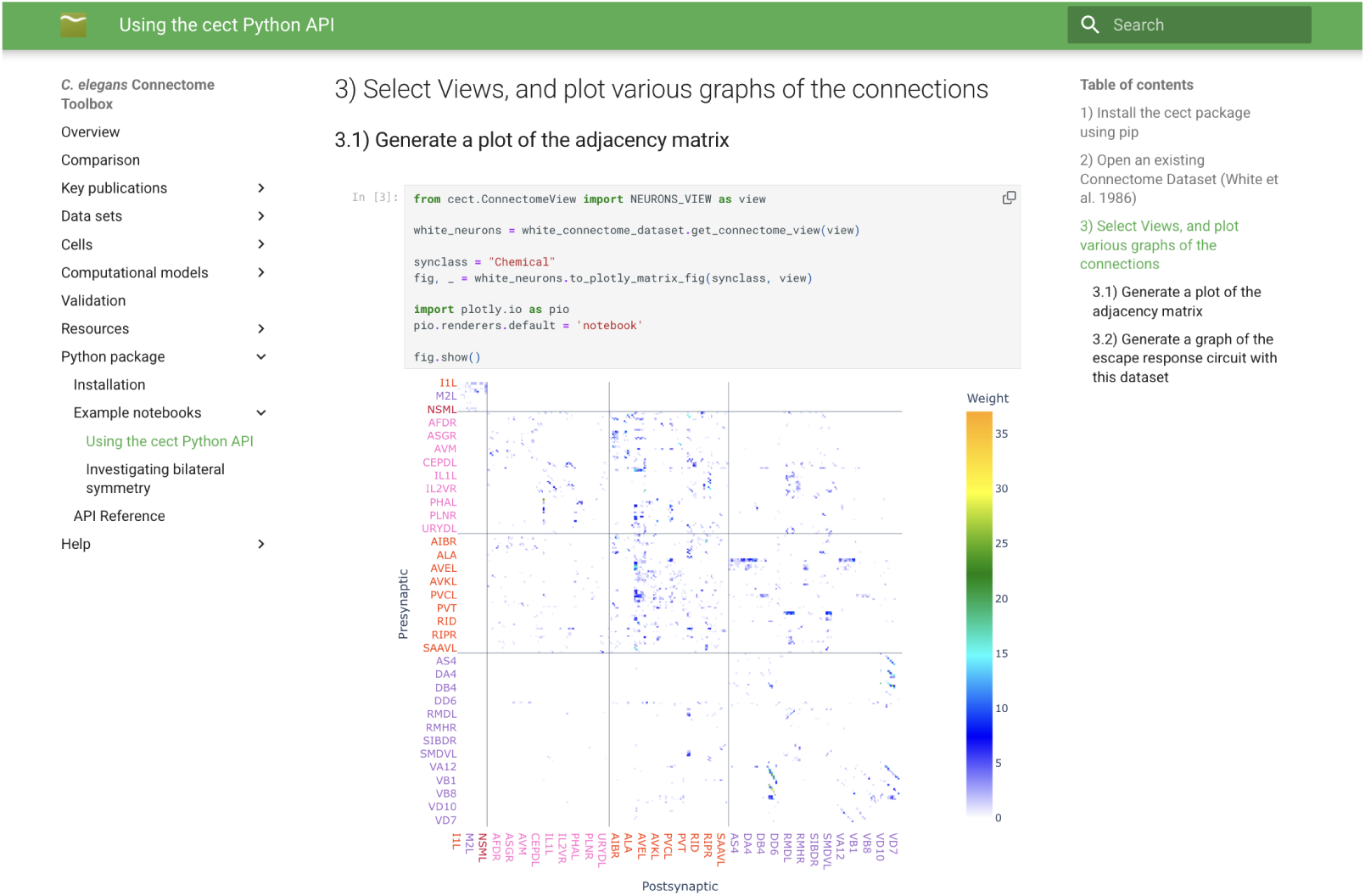
Screenshot of a section of a Jupyter notebook illustrating the use of the cect Python API. This illustrates how the interactive Plotly based graphs used on the pre-built website pages can also be generated locally in notebooks by users for their own analyses.

**Figure 3–Figure supplement 1.**
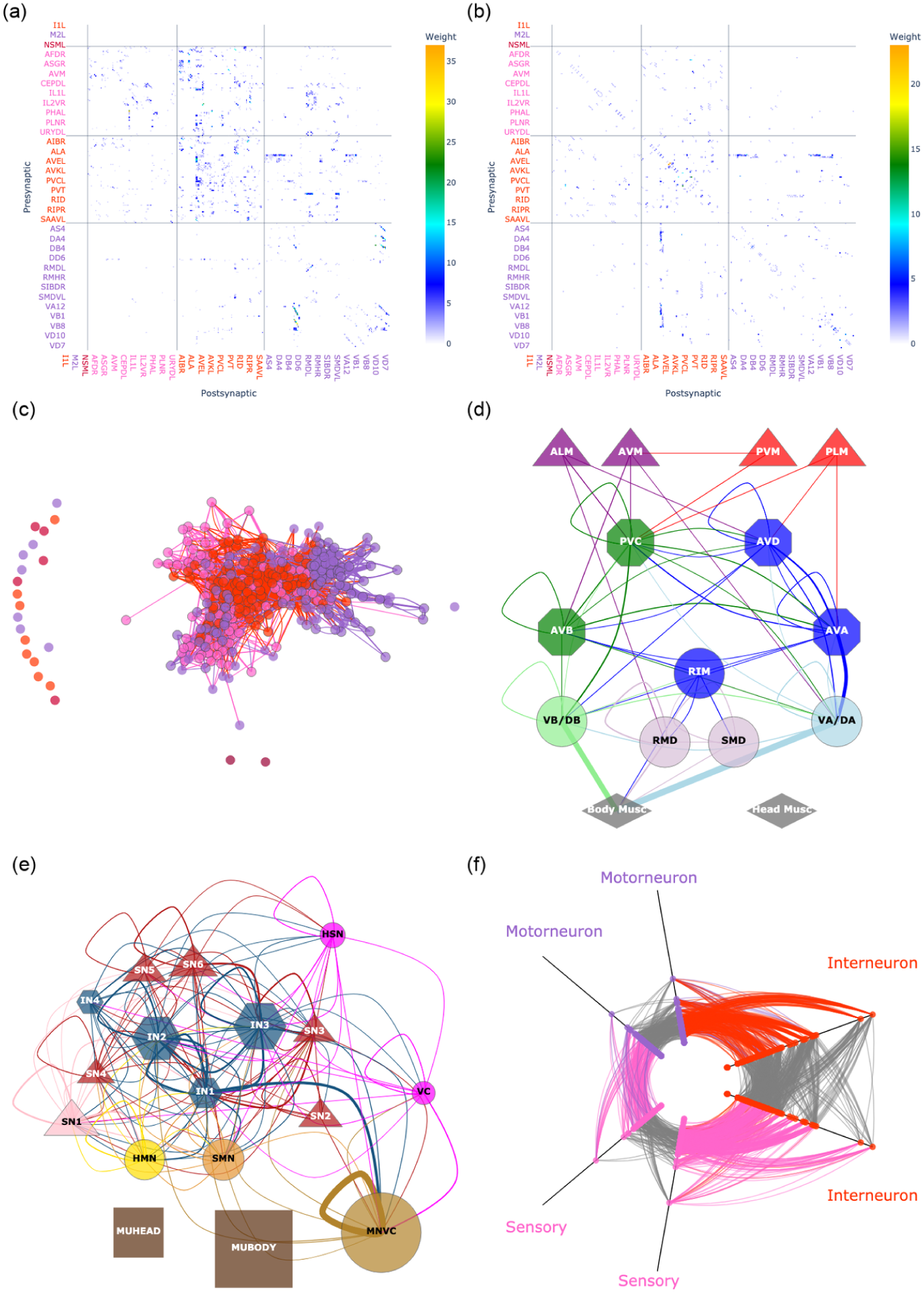
Different types of visualizations available in Connectome Toolbox, all generated from the same dataset, ***Varshney et al. (2011)***. (a) Matrix view showing chemical synapse connectivity. (b) Matrix view showing electrical synapse connectivity. (c) Force-directed graph view showing all neurons and their chemical connections. Connections to pharyngeal neurons are not present in the ***Varshney et al. (2011)*** dataset, and these cells are in a disconnected group on the left. (d) Escape response circuit view based on ***Pirri and Alkema (2012)***. The majority of nodes consist of the left and right neurons of a bilateral pair (e.g. ALML and ALMR), apart from single cell nodes (AVM, PVM), motorneuron nodes (VB/DB contains VB1, VB2, DB1, etc.; SMD contains SMDDL, SMDDR, SMDVL and SMDVR) and head/body muscle nodes (28, 67 muscles respectively). Note, in the original ***Varshney et al. (2011)*** dataset, all connections from motorneurons to muscles (i.e. neuromuscular junctions) are listed as connecting to a single endpoint, “NMJ”. This generic muscle is included in the body wall muscle group above to illustrate these connections. (e) Whole network view of chemical connections based on Fig. 3 in ***Cook et al. (2019)***. (f) Hive plot showing chemical connections between sensory neurons (pink), interneurons (red) and motorneurons (purple). Each of these are screenshots from the online interactive versions of the plots, which allow users to zoom in and hover over nodes/edges to identify specific cells and connections, e.g. the 2 red nodes representing highly connected interneurons at the end of the right axes in (f) can be identified as AVAL and AVAR by hovering over these nodes, with other “rich club” neurons (***Towlson et al., 2013***), AVB, AVD, AVE, PVC just below them.

**Figure 3–Figure supplement 2.**
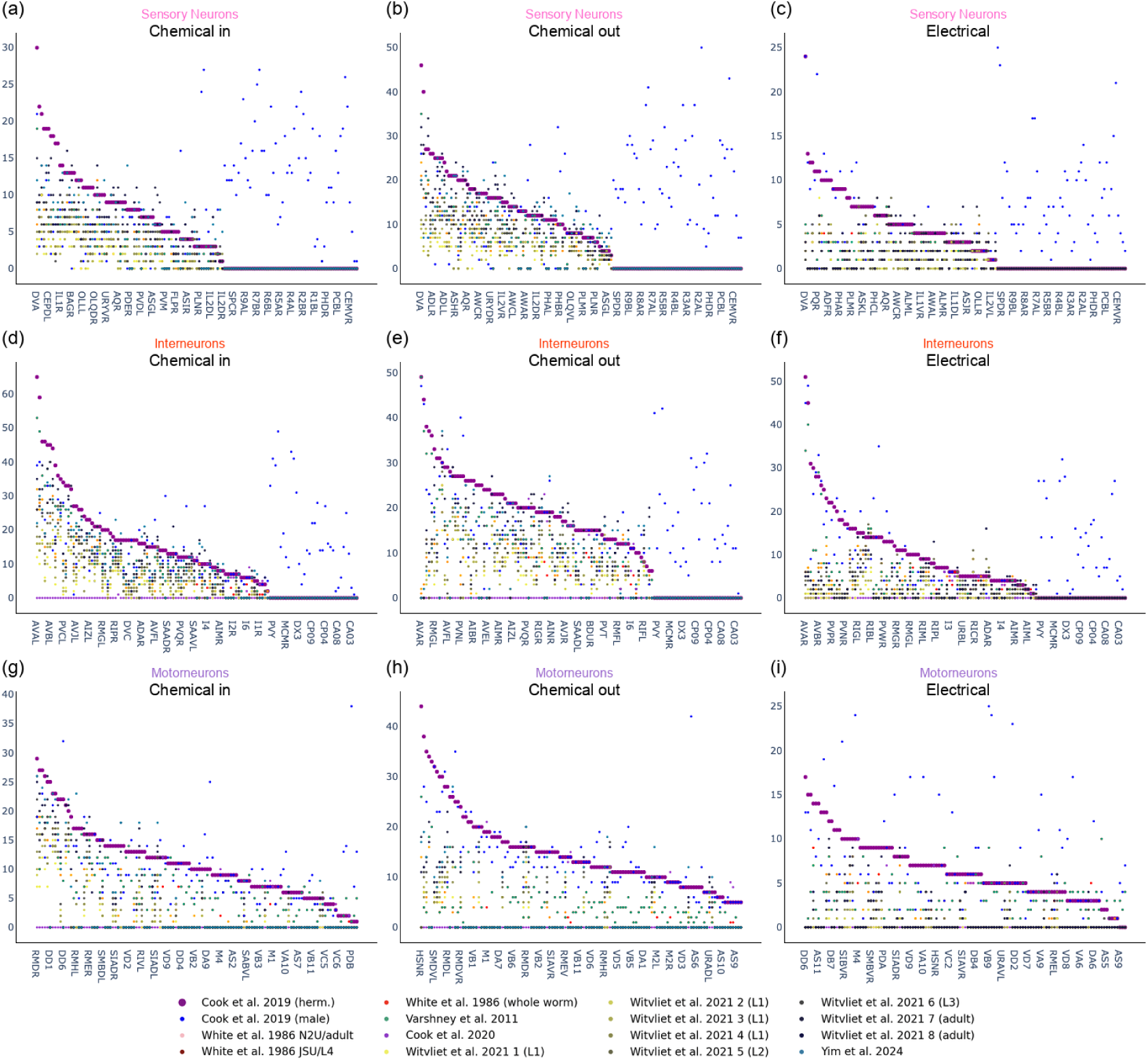
Comparison of numbers of connections to and from neurons across datasets. (a-c) Sensory neuron connections, showing the numbers of chemical connections coming into each neuron (a), chemical connections going out of each neuron (b), and the numbers of electrical connections (c). Neurons are ordered along the x-axis by the number of connections in the ***Cook et al. (2019)*** hermaphrodite dataset (which is plotted with larger, purple points), with other datasets aligned to this order for comparison (note, male only neurons are positioned on the right). As with the adjacency matrix visualizations (Fig. 4), not all cells are listed on the x-axis for clarity, but zooming in on the online interactive graph (or hovering over the points), allows identification of the cells (e.g. https://openworm.org/ConnectomeToolbox/Cells/#sensory-neuron). (d-f) Same as (a-c) but for interneurons. (g-i) Same as (a-c) but for motorneurons. A key for the colors of points used for datasets shown in (a-i) is given at the bottom of the figure.

**Figure 3–Figure supplement 3.**
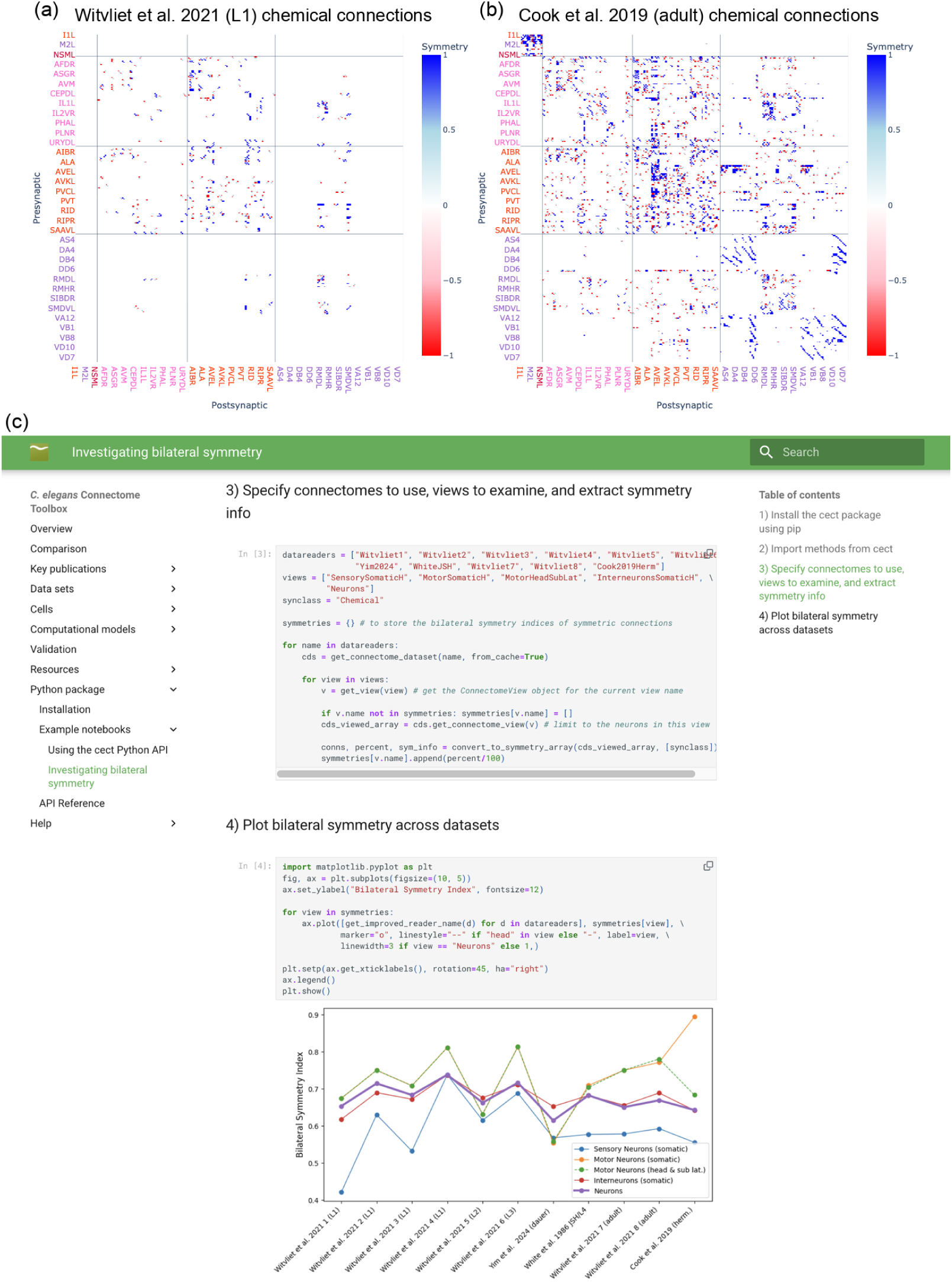
Analysis of the bilateral symmetry of connections across datasets. (a) Symmetry matrix for the chemical connections between neurons in the L1 larval dataset from ***Witvliet et al. (2021)***. This view is generated for all datasets online, and shows where bilaterally symmetrical connections (as defined in ***Kim et al. (2026)***) occur, or are absent. Bilaterally symmetrical connections (e.g. ADLL→ASEL and ADLR→ASER, or DVC→AVAL and DVC→AVAR) are shown by a dark blue square (symmetry value 1). Other connections which exist are shown by light blue (value 0.5), and the bilateral equivalent of each of these which is missing is shown in red (value −1). (b) Symmetry matrix for the chemical connections between neurons in adult hermaphrodite dataset from ***Cook et al. (2019)***. (c) Jupyter notebook illustrating how the Python API can be used to load multiple connectome datasets, apply different views, e.g. sensory or motorneurons only, and generate the symmetry arrays from each of these. The fraction of bilaterally symmetrical connections in each dataset can then be calculated from these arrays and plotted across datasets.

**Figure 6–Figure supplement 1.**
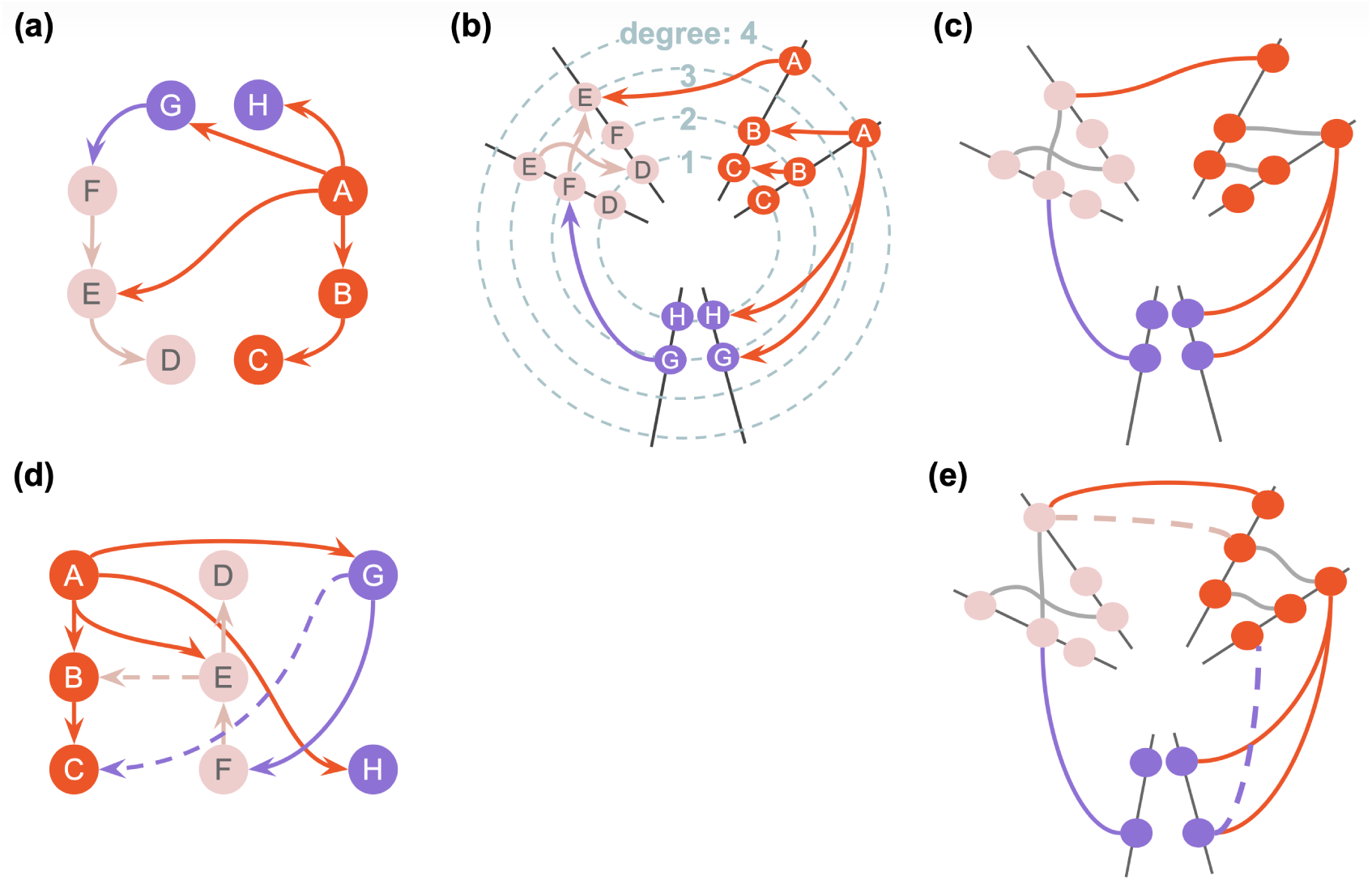
(a) A small network consisting of directed connections between 8 nodes of 3 different types (pink, red and purple). The layout of this network is non-deterministic and it can be displayed in various ways. (b) and (c) show the same network as (a) in the form of hive plots, with additional annotations in (b) (not normally shown) to facilitate interpretation. A hive plot (***Krzywinski et al., 2012***) is an alternative type of data visualization used to represent complex networks. Hive plots are composed of three main components: axes, nodes (arranged along the axes), and edges linking the nodes. The axes represent specific attributes of the node (often the degree of connectivity) and are arranged radially around a central point. In the case of our hive plots as shown in Fig. 6, the axes represent the 3 types of neurons, specifically sensory neurons, interneurons and motorneurons. While some hive plots have 3 axes, ours have 6 (2 for each type of neuron - straight black lines), allowing both connections to the other neuron types to be shown, as well as internal connectivity within a type. The positioning of the nodes along the axes is determined by the total degree of the node (sum of incoming and outgoing connections). To give examples of two connections in (b), red node A connects (with a red line indicating the source) to both purple node H (which is that neuron’s only connection) as well as purple node G, one of that node’s two connections, hence G (having degree 2) is further out radially along the axis than H (degree 1). Both of these purple nodes are depicted along the right hand (connections to/from red) and the left hand (connections to/from pink) axis, with one connection out of node G to pink node F on the left. (c) shows the same network as (b), but with labels, degrees and arrows removed (as is used in Fig. 6d-i). Edge coloring signifies the source node type, except for internal connections, which are all gray. The high level properties of the network are clear from this representation: internal connections are present in the pink and red groups, there are none in the purple; connections are present between all groups, but there are more from red to purple. (d) shows the same network as (a), but with 2 extra connections (dashed lines), and a different layout. It is not immediately clear that the majority of the connections between these two networks are the same. The similarities and differences between (a) and (d) are much easier to see by comparing the hive plot representations (c) and (e), with the new connections shown as dashed lines in (e).

**Figure 7–Figure supplement 1.**
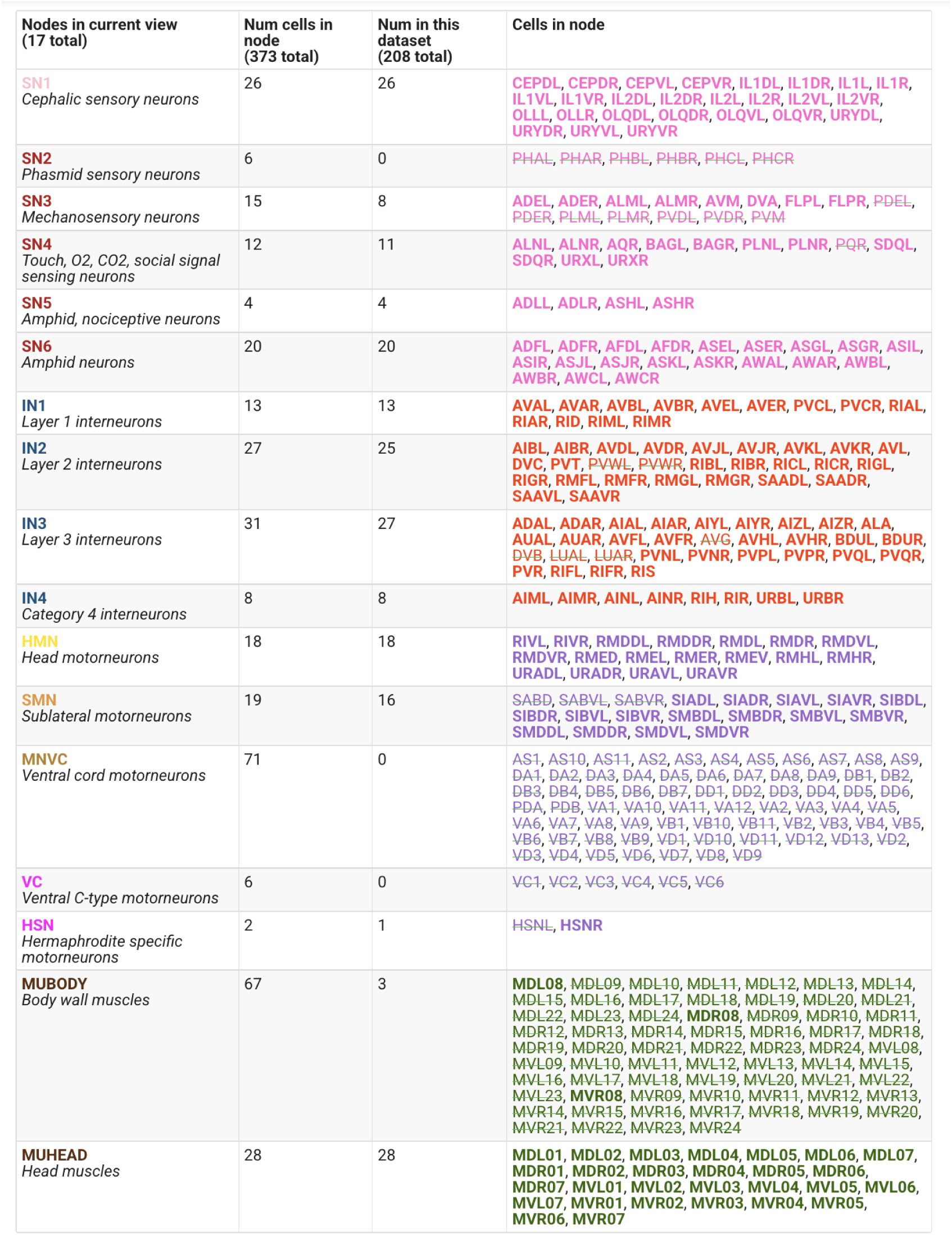
Table listing the specific cells present in each of the nodes in the ConnectomeView as used in the graphs of Fig. 7, along with a short description of each category. Colors in column 1 match node colors used in that figure. This is a screenshot from the *C. elegans* Connectome Toolbox web interface’s information panel for the view. It is specifically from the view of the ***Yim et al. (2024)*** dataset (dauer connectome; https://openworm.org/ConnectomeToolbox/Full1_Yim2024_data/#view-info), which does not include all neurons or muscles from the ***Cook et al. (2019)*** dataset. Included cells are listed in bold in the right hand column, missing cells are struck through (e.g. the phasmid sensory neurons in SN2 are not present in this dauer connectome).

**Figure 7–Figure supplement 2.**
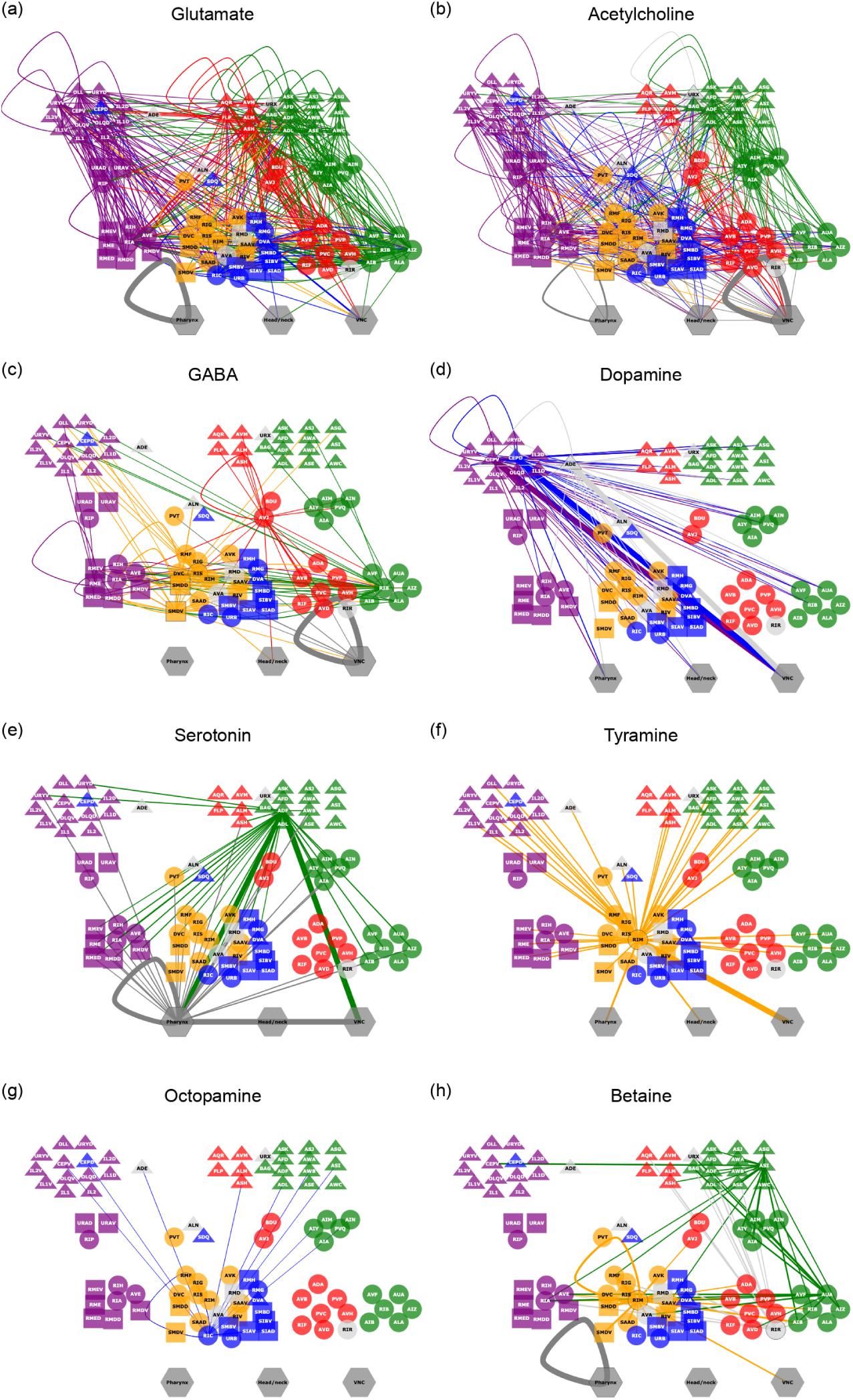
Comparing targets of neurotransmitters from ***Wang et al. (2024)*** overlaid on anatomical connectome from ***Cook et al. (2019)***, using Brainmap view of ***Brittin et al. (2021)***. This representation is based on the finding in that paper that the connectome is organized into five spatially ordered modules that support structured patterns of synaptic connectivity (anterior (purple nodes above), lateral (yellow), sublateral (blue), avoidance (red) and taxis (green); unclassified are light gray). These modules are arranged within a three-layer architecture in which sensory inputs feed forward through integration layers and converge onto a distributed, recurrent layer that coordinates motor output across the brain. Sensory neurons are represented as triangles, interneurons as circles, and motorneurons as rectangles. Hexagonal dark gray nodes represent: all cells of the pharynx; motorneurons of the head and neck; and ventral nerve cord motorneurons. No connections to muscles are included in this view. (a) Distribution of glutamatergic connections. (b) Acetylcholine. (c) GABA. (d) Dopamine. (e) Serotonin. (f) Tyramine. (g) Octopamine. (h) Betaine. An interactive example of one of these views can be seen online at https://openworm.org/ConnectomeToolbox/Brainmap_Wang2024Herm_data_graph/#gaba, which will indicate the cells present in each node.

**Figure 8–Figure supplement 1.**
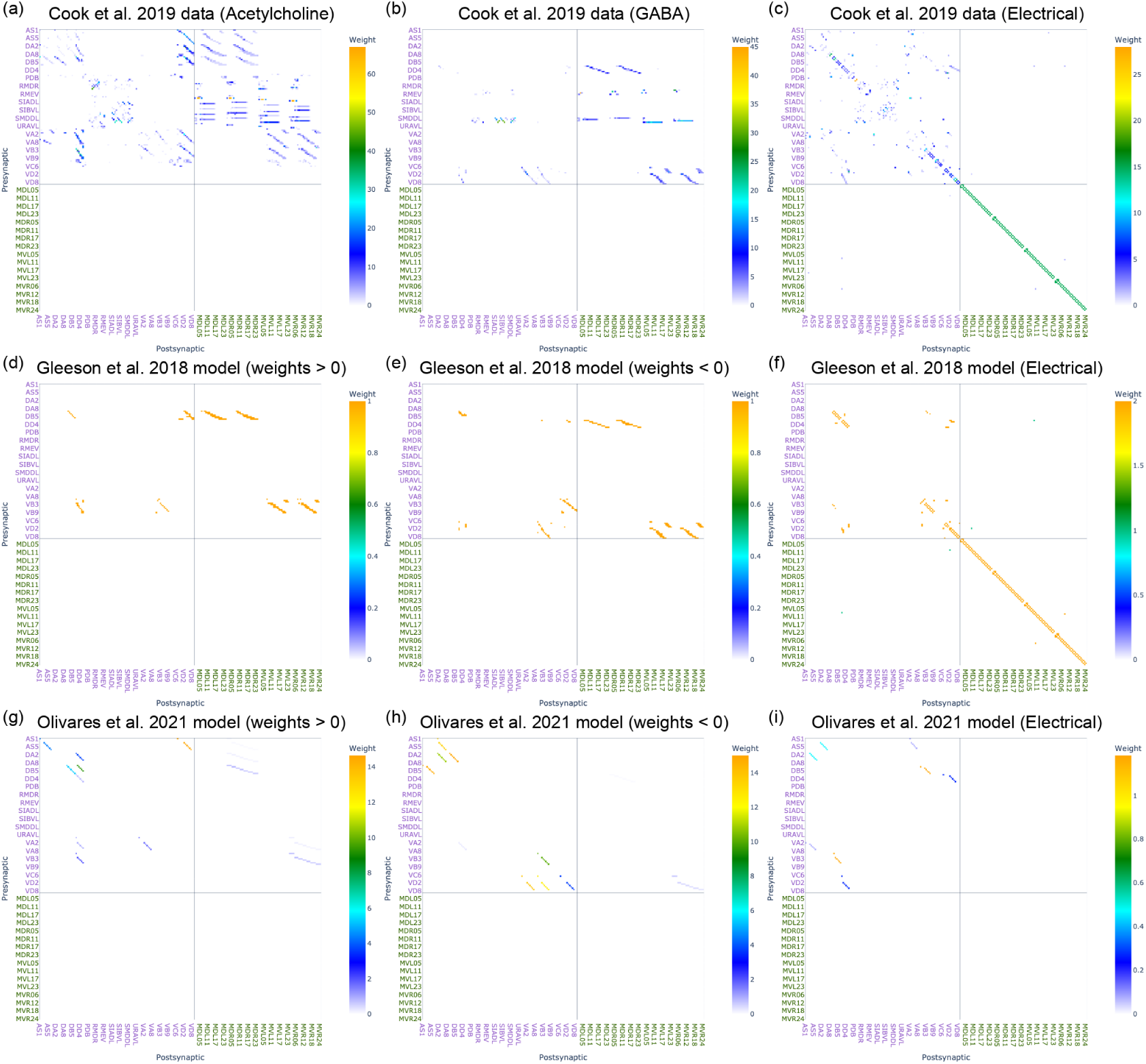
Adjacency matrix views of connectivity between motorneurons and muscles based on experimental data (***Cook et al., 2019***; ***Wang et al., 2024***) and computational models (***Gleeson et al., 2018***; ***Olivares et al., 2021***). All hermaphrodite motorneurons and muscles are included in the matrices, with the same ordering of cells in each panel, to facilitate comparison. Note that not all motorneurons are present in the two computational model networks. (a-i) Adjacency matrix views for the corresponding networks shown in Fig. 8(a-i). (a-c) Anatomical connectivity from ***Cook et al. (2019)***, coupled with neurotransmitter identities from ***Wang et al. (2024)***. (a) All chemical connections from cells releasing acetylcholine (putatively excitatory) according to ***Wang et al. (2024)***. (b) All chemical connections from cells releasing GABA (putatively inhibitory) according to ***Wang et al. (2024)***. (c) All electrical connections. (d-f) The connectivity used in the ***Gleeson et al. (2018)*** computational model. (d) All chemical connections with a positive weight, (e) with a negative weight, and (f) all electrical connections. (g-i) The connectivity used in the ***Olivares et al. (2021)*** computational model. (g) All chemical connections with a positive weight, (h) with a negative weight, and (i) all electrical connections.

## Notes

### Competing Interest Statement

The authors have declared no competing interest.

